# Alpha-synuclein and neuromelanin converge to promote Parkinson’s disease pathology

**DOI:** 10.64898/2026.10.19.760151

**Authors:** Alba Nicolau-Vera, Marta González-Sepúlveda, Joana M Cladera-Sastre, Thais Cuadros, Annabelle Parent, Gerard Roch, Jacopo Marongiu, Benjamin Huynh, Anahid Ansari Mahabadian, Feifei Su, Zac Chatterton, Javier Hoyo, Ariadna Laguna, YuHong Fu, Glenda Halliday, Miquel Vila

**Affiliations:** Vall d’Hebron Research Institute (VHIR)-Center for Networked Biomedical Research on Neurodegenerative Diseases (CIBERNED), 08035 Barcelona, Spain; Aligning Science Across Parkinson’s (ASAP) Collaborative Research Network, Chevy Chase, MD 20815, USA; Autonomous University of Barcelona (UAB), 08193 Cerdanyola del Vallès, Barcelona, Spain; The University of Sydney, Brain and Mind Centre & Faculty of Medicine and Health School of Medical Sciences, Sydney, NSW 2050, Australia; Catalan Institution for Research and Advanced Studies (ICREA), 08035 Barcelona, Spain

**Keywords:** Parkinson’s disease, alpha-synuclein, neuromelanin, dopamine, substantia nigra

## Abstract

Parkinson’s disease (PD) is defined by the preferential degeneration of neuromelanin (NM)-containing dopamine-producing neurons in the substantia nigra (SN), accompanied by intraneuronal inclusions rich in alpha-synuclein (αSYN). However, the mechanistic relationship between NM accumulation and αSYN pathology remains unclear. Here, we reveal a bidirectional pathogenic relationship between αSYN and NM. In human post-mortem midbrain tissue, ventral nigral neurons with high NM content display increased αSYN mRNA expression and are preferentially lost in PD. In a rat model that recapitulates age-dependent, human-like NM accumulation via targeted tyrosinase (TYR) expression, co-expression of αSYN accelerates NM buildup, enhances αSYN oligomerization, and aggravates nigrostriatal degeneration. Mechanistically, αSYN oligomerization in NM-producing rats decreases vesicular dopamine storage and impairs vesicle fusion with the plasma membrane, increasing cytosolic dopamine availability for oxidation into NM. Together, these findings support a pathogenic feedback loop in which elevated αSYN accelerates NM synthesis through disrupted dopamine compartmentalization, while NM accumulation further promotes αSYN aggregation and neuronal loss. This αSYN-NM axis may underlie the preferential vulnerability of pigmented nigral neurons in PD and provide a mechanistic framework for the development of future targeted therapeutics.

## Introduction

Alpha-synuclein (αSYN) and neuromelanin (NM) are two central components in the neuropathologic definition of Parkinson’s disease (PD) (1,2). The definitive post-mortem diagnosis of PD relies on the loss of NM-laden neurons in the substantia nigra pars compacta (SNpc) and the presence of abnormal αSYN aggregates forming intracytoplasmic Lewy body (LB) inclusions (2). Together, these features represent the fundamental pathological signatures of PD, linking molecular protein aggregation with selective neuronal vulnerability and degeneration.

NM is a dark, insoluble pigment that progressively accumulates within specific catecholaminergic neurons of the human brain, most notably in the SNpc, starting in early childhood (3,4). These neurons, rich in the neurotransmitter dopamine, gradually become visibly pigmented as NM deposits build up over time (5). Although the precise physiological role of NM, if any, remains incompletely understood, it is thought to participate in cellular homeostasis by sequestering potentially toxic catecholamine metabolites and metals such as iron (6,7). Notably, the same population of NM-laden neurons is among the earliest and most severely affected in PD (5). This spatial and temporal correlation between NM accumulation and neuronal vulnerability has led to the hypothesis that NM may influence susceptibility to neurodegeneration, either by modulating oxidative stress (8), interfering with metal homeostasis (7,9), disrupting neuronal proteostasis (10–12), or through other yet unidentified mechanisms (13). Consequently, the study of NM dynamics in the SN offers critical insights into both normal dopaminergic neuron physiology and the pathogenesis of PD.

In humans, classical nigral LB, which are a hallmark of PD, and their presumed precursor structures, pale bodies, typically form within the intracellular areas of the neuronal cytoplasm in which NM accumulates, often developing in close physical association with this pigment (14,15). αSYN redistributes to the NM pigment in early stages of PD and becomes entrapped within NM granules (16,17). Additionally, there is an age-related accumulation of αSYN protein within pigmented nigral neurons, even in the absence of clinical PD (18,19). Despite the established associations between NM, αSYN, and PD, the mechanistic relationship between αSYN and NM in the context of PD pathogenesis remains poorly understood. While αSYN has been extensively incorporated into experimental models of PD (20,21), NM is absent in most animal species, including rodents, and is therefore typically not accounted for in current *in vivo* experimental paradigms (22). To overcome this major limitation, we recently developed rodent models exhibiting age-dependent production and accumulation of human-like NM within PD-vulnerable neurons, at levels up to those reached in elderly humans (12,23–25). These models are based on the expression of human melanin-producing enzyme tyrosinase (TYR) in the brain (12,23–25). Experimentally inducing NM production in TYR-expressing rodents and non-human primates stimulates LB-like inclusion formation and neurodegeneration (12,23–25). To investigate the relationship between αSYN and NM, we employed an *in vivo* model combining αSYN overexpression with TYR-induced NM production. This strategy enabled us to determine whether elevated αSYN levels, as observed in PD, influence the extent and temporal progression of NM-associated PD pathology. Moreover, it provides a framework to explore the molecular mechanisms linking αSYN aggregation, age-dependent pigment accumulation, and neuronal vulnerability in PD.

## Materials and methods

### Post-mortem human midbrain tissue samples

Tissue sections from pathologically confirmed cases of PD (n=14) (26) and controls without neurological or neuropathological disease (n=18) were obtained from the Sydney Brain Bank following project-specific ethics approval (University of Sydney Human Research Ethics Committee 2021/HE000845). All cases with PD were levodopa-responsive and fulfilled the UK Brain Bank Clinical Criteria for a diagnosis of clinical PD (27). Two PD cases have concomitant intermediate Alzheimer’s disease-like neuropathological changes (28). The demographic details for the groups are summarized in **Table 1**.

**Table 1.** Group demographics (mean ± standard deviation)

|  | Controls for GeoMx | Controls for IHC | PD cases for IHC |
|---|---|---|---|
| Sex (M/F) | 3/7 | 4/4 | 10/4 |
| Age at death (y) | 90 ± 7 | 81 ± 10 | 76 ± 5 |
| PM delay (hr) | 28 ± 14.5 | 24.5 ± 13.5 | 19 ± 15 |
| Braak LP stage | 0 ± 0 | 0 ± 0 | 5.1 ± 0.7<sup>***</sup> |
| A score (28) | 1.3 ± 0.8 | 0.8 ± 0.8 | 0.8 ± 1.1 |
| B score (28) | 1.1 ± 0.7 | 1.0 ± 0.8 | 1.1 ± 0.9 |
| C score (28) | 0.6 ± 1.0 | 0.5 ± 1.1 | 0.4 ± 0.7 |

*Comparisons were made with univariate tests. For the gender difference, the chi-square test compared to IHC controls was applied, p < 0.001<sup>\*\*\*</sup>. A=regional distribution of amyloid-β deposits, B=regional distribution of neurofibrillary tangles, and C=semiquantitative density of cortical neuritic plaques; together, these scores are used to determine the level of Alzheimer disease neuropathologic change. M=male, F=female, IHC=immunohistochemistry, LP=Lewy pathology, PM=post-mortem.*

### Immunohistochemistry, imaging, and quantifying human dopamine neurons

Formalin-fixed, paraffin-embedded 8 μm transverse midbrain sections at the level of the exiting third nerve were immunostained with an antibody for TH (mouse monoclonal, Sigma T2928, diluted 1:200) following antigen retrieval with sodium citrate buffer (pH 6.0) in a programmable antigen retrieval cooker (Aptum Bio Retriever 2100, Aptum Biologics Ltd, UK). Primary antibody binding to the epitope was visualised using SignalStain® Boost IHC Detection Reagent (HRP, Mouse) (Cell Signaling, #8125) and Black DAB (stable) substrate/chromogen 2-component system (INNOVEX bioscience, #NB319-BB70) according to the manufacturer’s instructions. DPX mounting media (Sigma-Aldrich, Cat. No. 317616) was used to coverslip sections.

The entire TH immunoreactive region in each midbrain section was scanned using an Olympus VS120 Slide Scanner at 20x magnification with auto-focus and auto-exposure settings. The images were processed using VS-desktop software (Olympus Soft Imaging Solutions GmbH, version 2.9 13753) and converted to TIFF format. Any subsequent image contrast adjustments performed in ImageJ (National Institutes of Health, USA, version 1.52e) were used exclusively for figure presentation and were not part of the quantitative analysis pipeline.

The ventral and dorsal tiers of the SNpc and the ventral tegmental area (VTA) were identified using the free-hand ROI tool on QuPath (version 0.3.2), and their areas were calculated. As previously described (29,30), dopamine neurons were categorized into distinct subpopulations based on TH-immunoreactivity (grey) and NM signal (brown) as TH^+^NM^-^, TH^+^NM^+^, and TH^-^ NM^+^. Cells were classified as TH^-^ when TH immunoreactivity was not readily identifiable during systematic cell counting. These dopamine neuron types were identified across different ROIs and counted manually using the point annotation tool in QuPath (version 0.3.2), and their densities were calculated from ROI areas. Repeat counts on 10% of the sections varied by <3%, and interrater variations were <4%. The resulting density of dopamine neuron types per region was used in the downstream statistical analyses.

### Gene expression profiling of human dopamine neurons

Formalin-fixed, paraffin-embedded 6-μm transverse midbrain sections at the level of the exiting third nerve were used for GeoMx. Tissue sections were stained with Alexa Fluor® 647 anti-TH (BioLegend 818008) following the manufacturer’s instructions (GeoMx® DSP Manual Slide Preparation, version MAN-10150-03).

The ventral and dorsal tiers of the SNpc and the VTA were identified and ROIs (500 x 600 μm^2^) within those regions processed using the Nanostring GeoMx® Digital Spatial Profiler, according to the manufacturer’s instructions. In total, 49 ROIs from the ventral tier of the SNpc (SNv) and 80 ROIs from the dorsal tier of the SNpc (SNd) and VTA were analyzed. Sequencing of retrieved mRNA probes yielded spatial transcriptomic profiles for 18,675 transcripts, of which 9,012 exceeded the limit of quantification and were retained for downstream analyses. Counts were used for statistical analysis and normalized gene expression (described in (31)) were used for plotting.

### Animals

Adult male Sprague-Dawley rats (Charles River), 225–250 g and 8 weeks of age at the time of surgery, were housed two or three per cage with *ad libitum* access to water and food and exposed to a 12 hours (h) light/dark cycle. All experimental and surgical procedures were conducted in strict accordance with the European Directive (2010/63/UE) and Spanish regulations (Real Decreto 53/2013; Generalitat de Catalunya Decret 214/97) on the protection of animals used for experimental and other scientific purposes. Protocols were approved by the Vall d’Hebron Research Institute (VHIR) Ethical Experimentation Committee and the Generalitat de Catalunya (Protocol 11442). Rats were randomly distributed into the different experimental conditions and experimental groups were processed at the same time to minimize bias.

### Viral vector production

rAAV vector serotype 5 expressing human αSYN cDNA driven by the chicken beta-actin (CBA) promoter (AAV5-CBA-hαSYN; concentration 1×10^13^ gc/ml) was obtained at the Michael J. Fox Foundation. rAAV serotype 2/1 expressing human TYR cDNA driven by the cytomegalovirus (CMV) promoter (AAV2/1-CMV-hTYR) was produced at the Viral Vector Production Unit of the Autonomous University of Barcelona (UPV-UAB, Spain). Briefly, rAAV vector expressing hTYR was produced by triple transfection of 2 × 10^8^ HEK293 cells with 250 μg of pAAV, 250 μg of pRepCap, and 500 μg of pXX6 plasmid mixed with polyethylenimine (PEI; branched, MW 25000; Sigma-Aldrich). The UPV-UAB generated a pAAV plasmid containing the ITRs of the AAV2 genome, a multicloning site to facilitate cloning of expression cassettes, and ampicillin resistance gene for selection. 48 h after transfection, cells were harvested by centrifugation [200 × g, 10 minute (min)]; resuspended in 30 ml of 20 mM NaCl, 2 mM MgCl_2_, and 50 mM Tris-HCl (pH 8.5), and lysed by three freeze–thaw cycles. Cell lysate was clarified by centrifugation (2000 × g, 10 min) and the rAAV particles were purified from the supernatant by iodixanol gradient as previously described (32). The clarified lysate was treated with 50 U/ml of Benzonase (Novagen; 1 h, 37 °C) and centrifuged (3000 × g, 20 min). The vector-containing supernatant was collected and adjusted to 200 mM NaCl using a 5 M stock solution. To precipitate the virus from the clarified cell lysate, polyethylene glycol (Sigma-Aldrich, #PEG8000) was added to a final concentration of 8% and the mixture was incubated (3 h, 4 °C) and centrifuged (8000 × g, 15 min). The AAV containing pellets were resuspended in 20 mM NaCl, 2 mM MgCl_2_, and 50 mM Tris-HCl (pH 8.5) and incubated for 48 h at 4 °C. The rAAV titration method used was based on the quantitation of encapsulated DNA with the fluorescent dye PicoGreen® as previously described (33), and the following vector concentration was obtained: AAV-hTYR: 2.43×10^12^ gc/ml.

### Stereotaxic surgery

Before surgery, all surgical instruments were autoclaved and analgesic solution (meloxicam, 0.5 mg/ml) was prepared. All surgical procedures were performed with the animals placed in stereotaxic frame under general anaesthesia with isoflurane (Baxter) (5% for the induction phase and 2% for the maintenance phase). Vector solutions were injected using a 10 µl Hamilton syringe fitted with a glass capillary (Hamilton, #701). In all cases, infusion was performed at a rate of 0.4 µl/min, with the needle remaining in place for an additional 4 min before being slowly retracted. Injection was performed unilaterally on the right side of the brain at the corresponding coordinates relative to bregma according to the stereotaxic atlas of Paxinos and Watson (34) and the craniotomy site was sealed by suturing the scalp with sterile sutures. Rats received 2 µl of a 1:1 mixture of either AAV-αSYN/vehicle, AAV-TYR/vehicle or AAV-αSYN/AAV-TYR above the SNpc: antero-posterior: −5.2 mm, medio-lateral: −2 mm, dorso-ventral: −7.6 mm (Supplementary Fig. 1). AAV5-CBA-hαSYN was administered at 1×10¹³ gc/ml, corresponding to 1×10¹□ gc per injection, and AAV-hTYR at 2.43×10¹² gc/ml, corresponding to 2.43×10□ gc per injection. To reduce pain and inflammation, meloxicam (5 mg/kg) was administered subcutaneously. Because previous studies using the same nigral TYR paradigm showed that sham injections and empty-vector AAV controls do not induce NM accumulation or the nigrostriatal pathology elicited by TYR expression (12), these additional control groups were not included in the present study in order to reduce animal numbers in accordance with the Reduction principle of the 3Rs. Instead, the contralateral non-injected hemisphere of each animal served as an internal reference.

**Supplementary Figure 1.**
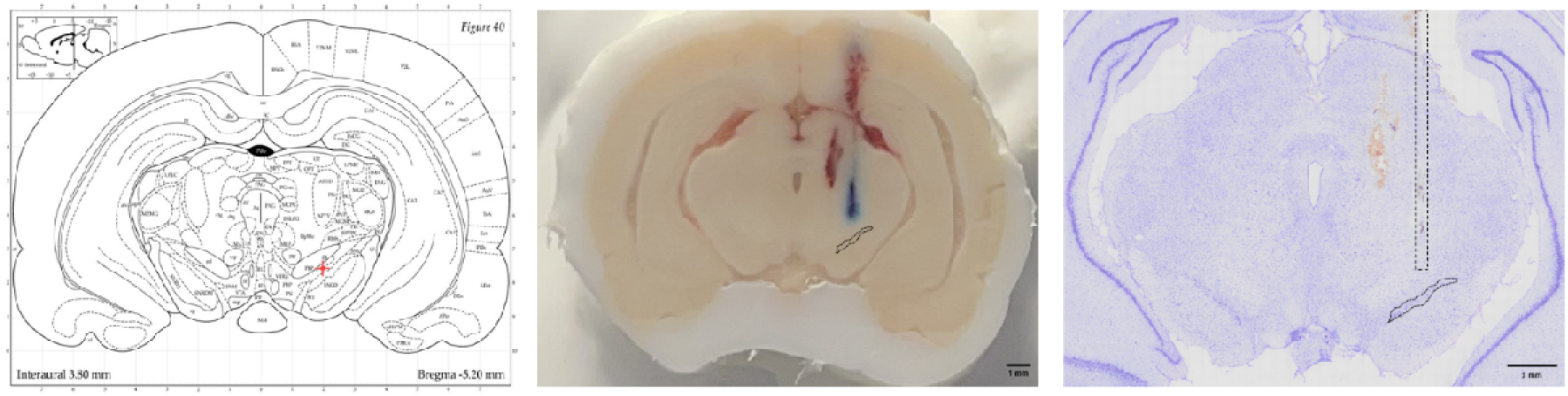
Anatomical localization of the infusion site and needle track. **Left**, coronal plate from the Paxinos and Watson rat brain atlas (1982), with the red cross marking the infusion site immediately above the substantia nigra pars compacta (SNpc) to minimize direct mechanical damage to this region (AP, −5.2 mm; ML, −2.0 mm; DV, −7.6 mm relative to bregma). Cresyl violet was infused unilaterally into the right hemisphere at 0.4 μL/min, with the needle left in place for 3 min before withdrawal. **Middle**, coronal view during cryostat sectioning, showing cresyl violet deposition (blue) above the SNpc (dashed contour). The rat was perfused under deep isoflurane anesthesia 6 h after infusion, and the brain was post-fixed, cryoprotected, and frozen. **Right**, corresponding 20-μm Nissl-stained coronal section from the same animal, showing the needle track (dashed rectangle) and the SNpc (dashed contour). Scale bars: 1 mm.

### Brain processing

Animals were deeply anesthetized with isoflurane and then perfused through the left ventricle with saline (0.9% [wt/vol]) at room temperature (RT) for 5 min followed by 4% paraformaldehyde fixative solution in phosphate buffer (Panreac, #252931.1214) for 15 min, before brains were removed for histology. Brains were post-fixed for 24 h in the same fixative solution. After fixation, rat brains were washed with PBS and processed for paraffin embedding at the VHIR’s Drug Delivery and Targeting laboratory facility. Paraffin-embedded rat brains were sectioned with a slicing microtome (Leica, Germany) at 5 µm-thickness and sections containing the SN and striatal (STR) regions were collected in Polysine adhesion slides (Epredia^TM^, #J2800AMNZ).

### Immunostaining

Paraffin-embedded brain sections were first deparaffinized and rehydrated. Slides were incubated at 60 °C for 30 min to melt the paraffin, followed by three washes in xylene (3 min each) to remove residual paraffin. Rehydration was performed through a descending ethanol gradient (100%, 95%, and 70%; 10 min each), after which slides were washed twice in buffer (0.1 M Tris-buffered saline [TBS] for immunohistochemistry or 0.1 M phosphate-buffered saline [PBS] for immunofluorescence).

For immunohistochemistry, endogenous peroxidase activity was blocked by incubating sections in 3% H□O□ and 10% methanol in 0.1 M TBS for 10 min, followed by three washes in TBS (5 min each).

Antigen retrieval was performed by incubating slides in 10 mM citrate buffer (pH 6.0) at 95 °C for 20 min, followed by cooling at room temperature for 20 min. Sections were then permeabilized with two washes (5 min each) in buffer containing 0.5% Triton X-100, followed by a final wash in buffer alone. Slides were placed in a humid chamber, excess liquid was removed, and sections were outlined using a hydrophobic pen.

Non-specific binding was blocked by incubating sections with 5% normal goat serum (NGS) in buffer for 1 h at room temperature. Primary antibodies, diluted in 2% NGS in the corresponding buffer, were applied and incubated at 4 °C for 24–48 h (depending on the antibody). After incubation, sections were washed three times (5 min each) in buffer.

Detection was then performed according to the staining method. For immunohistochemistry, sections were incubated with biotinylated secondary antibodies (1 h, room temperature), followed by avidin-biotin complex (ABC) solution for 1 h. After washing, signal was developed using DAB (brown) or Vector® SG (dark blue) for 5-15 min, depending on the desired intensity. Sections were then washed, dehydrated through ascending ethanol concentrations (70%, 95%, 100%), cleared in xylene, and mounted with DPX mounting medium. Slides were allowed to dry overnight.

For immunofluorescence, sections were incubated with appropriate Alexa Fluor-conjugated secondary antibodies (e.g., 488, 568, 594, 647) together with Hoechst 33342 (1:5000) for nuclear staining (1 h, room temperature). After washing, slides were coverslipped using fluorescent mounting medium.

Lists of primary and secondary antibodies are provided in Tables 2 and 3.

**Table 2.** List of primary antibodies.

| Primary antibody | Host | Manufacturer | Technique | Brain region | Dilution | Incubation |
|---|---|---|---|---|---|---|
| Anti-αSYN | Rat | Enzo, #ALX-804-258-LC05 | IHC | SN, STR | 1:1000 | 24h |
|  |  |  | IF | SN | 1:100 | 24h |
| Anti-pαSYN | Rabbit | Abcam, #51253 | IHC | SN | 1:500 | 24h |
| Anti-CD68 | Mouse | BIO-RAD, #MCA341R | IHC | SN | 1:100 | 24h |
| Anti-p62 | Guinea pig | Progen, #GP62-C | IF | SN | 1:500 | 24h |
| Anti-TH | Rabbit | Calbiochem, #657012 | IHC | SN | 1:20,000 | 48h |
|  |  |  |  | STR | 1:3,500 | 48h |
|  |  |  | IF | SN | 1:1000 | 24h |
| Anti-TYR | Mouse | Epredia<sup>TM</sup>, #MS-800-P1 | IHC | SN | 1:750 | 24h |
| Anti-VAMP2 | Mouse | Thermo Fisher Scientific, #MA5-38260 | IF | SN | 1:500 | 24h |

*IHC, immunohistochemistry; IF, immunofluorescence.*

**Table 3.** List of secondary antibodies.

| Technique | Secondary antibody | Host | Manufacturer | Dilution |
|---|---|---|---|---|
| IHC | Biotinylated anti-rabbit | Goat | Vector laboratories, #BA1000 | 1:1000 |
|  | Biotinylated anti-mouse | Goat | Vector laboratories, #BA9200 | 1:1000 |
|  | Biotinylated anti-rat | Goat | Vector laboratories, #BA9400 | 1:1000 |
| IF | Alexa fluor 488 anti-guinea pig | Goat | Gibco, #A11073 | 1:1000 |
|  | Alexa fluor 488 anti-mouse | Goat | Gibco, #A11001 | 1:1000 |
|  | Alexa fluor 647 anti-rabbit | Donkey | Life technologies, #A31573 | 1:1000 |
|  | Alexa fluor 647 anti-rat | Chicken | Gibco, #A21472 | 1:1000 |

*IHC, immunohistochemistry; IF, immunofluorescence.*

### Proximity-ligation assay (PLA)

The day before the assay, rabbit anti-αSYN polyclonal antibody (Merck, #AB5038P) was conjugated with Duolink probes. 2 µl of Conjugation buffer from the Duolink PLUS (Merck, #DUO92009-1KT) or Duolink MINUS (Merck, #DUO92010-1KT) probes was added to 20 µL of rabbit anti-αSYN polyclonal antibody (Merck, AB5038P) at a concentration of 1 mg/ml. Antibody solution was transferred to one vial of lyophilized oligonucleotides (either PLUS or MINUS) and incubated at RT overnight. 2 µl of Stop Reagent was added to the reaction and incubated at RT for 30 min. Finally, 24 µl of Storage Solution was added, and the solution was stored at 4 °C.

For the assay, rat paraffin-embedded brain sections were initially deparaffinized and rehydrated. To unmask epitopes, sections were submerged in citrate buffer 10 mM, pH 6 and incubated in water bath at 95°C for 20 min followed by 20 min at RT. Slides were then washed 2 x 5 min in PBS 1x-triton 0.5% buffer followed by 1 x 5 min in PBS 1x buffer. For blocking, tissue sections were dried and circled with a hydrophobic pen (Vector laboratories, #NC9545623). Tissue sections were then incubated in Duolink blocking solution from Duolink PLUS or MINUS (1-2 drops/section). Conjugated primary antibodies (PLUS and MINUS) were diluted at 1:500 with the antibody diluent. Tissue sections were then incubated with the primary antibodies overnight at 4 °C. The next day, slides were washed 2 x 5 min in 1x Wash Buffer A (Merck, #DUO82049-4L). For ligation, 5x Duolink ligation buffer was diluted in 1:5 high purity water and then mixed to create the ligation buffer. Ligase was then diluted in ligation buffer at 1:40. Tissue sections were then incubated for 30 min at 37°C. Slides were washed 2 x 5 min in 1x Wash Buffer A (Merck, DUO82049-4L). For amplification, 5x Duolink amplification buffer was diluted in 1:5 high purity water and then mixed to create the amplification buffer, which is light-sensitive. Polymerase was then diluted in amplification buffer at 1:80. Tissue sections were then incubated 100 min at 37 °C. For final washes, slides were washed 2 x 10 min in 1x Wash Buffer B (Merck, #DUO82049-4L) and in 0.01x Wash B for 1 min. Tissue sections were then incubated with Hoechst 33342 (Thermo Fisher Scientific, #H3570) at 1:5000 for 10 min at RT for nuclei visualization. Slides were mounted with DAKO fluorescent mounting medium (Merck, #F4680) and stored at 4 °C.

The PLA immunopositive area was quantified in a 5 μm-thick section of the rat SN using QuPath software. Immunofluorescence-stained sections were scanned with the Olympus Slideview VS200 slide scanner and visualized using Olyvia 3.3 software. For quantification, the SNpc was selected using the ROI manager in QuPath. Positive cell detection analysis was performed with a pixel size of 0.5 μm and an intensity threshold of 100 within the Cy3 detection channel to determine the total immunopositive area.

### Neuropathological quantifications

For neuron cell counting, tyrosine hydroxylase (TH)-, αSYN- and phosphorylated αSYN (pαSyn)-immunostained sections were scanned with Olympus Slideview VS200 slide scanner and the Olyvia 3.3 software. For TH immunostaining, 5 µm-thick rat SNpc brain sections were collected every 17th section, yielding 10-12 sections per animal. For αSYN and pαSyn immunostaining, four representative 5 µm-thick rat brain sections spanning the entire SNpc were selected. A specific artificial intelligence (AI)-assisted algorithm was implemented for the quantification of Vector^®^SG-immunostained neurons (including TH-, αSYN- and pαSyn-positive neurons) using the Olympus VS200 Desktop 3.3 software. The algorithm was trained to identify four categories of objects within the desired region of interest (ROI): TH^+^NM^-^, TH^+^NM^+^, TH^-^NM^+^ neurons and extraneuronal NM. Briefly, the algorithm for the simultaneous classification and quantification of neuronal phenotypes and extraneuronal NM was trained using 28 representative images from TH-immunostained sections obtained from staining experiments performed on different days, thereby incorporating normal technical variability. These images were analyzed by multiple investigators to establish consistent annotation criteria. A total of 405 TH□NM□ neurons, 409 TH□NM□ neurons, 301 TH□NM□ neurons, 478 extraneuronal NM objects, and 817 background objects were manually annotated and used to train the algorithm. After 500,000 training iterations, the resulting algorithm achieved a similarity index of 0.70. Its performance was subsequently assessed on both the images used for training and additional images from the same staining experiments. Furthermore, all experimental outputs generated by the algorithm were manually reviewed, and any classification errors were corrected before the final quantitative analyses. Additionally, the algorithm was also capable to recognize αSYN and pαSyn-positive neurons.

For the quantification of TYR-immunopositive neurons, 5µm-thick rat paraffin-embedded SN sections immunostained for TYR were scanned using the Olympus Slideview VS200 slide scanner and the Olyvia 3.3 software, with one representative section selected per animal. TYR-positive neurons were manually quantified with QuPath and data was normalized to the total SN area to minimize bias.

For the quantification of intracellular NM levels, 5µm-thick rat paraffin-embedded hematoxylin and eosin (H&E)-stained sections containing SN at the same anatomical plane were selected (one representative section per animal), and all neurons per section were analysed. Catecholaminergic neurons were identified by the visualization of unstained NM brown pigment. H&E-stained sections were scanned using the Olympus Slideview VS200 slide scanner and the Olyvia 3.3 software. Quantifications were performed using QuPath for image visualization and ImageJ for analysis. Optical densitometry (OD) was used to assess intracellular NM density by measuring pixel intensity within NM-laden cells, excluding the nucleus, and correcting for nonspecific background, as previously described (12). Briefly, background was estimated from regions devoid of NM, and intracellular NM signal was calculated relative to this reference. Background-corrected OD values at or below zero were considered to indicate no detectable NM above background. Such values can arise from small local variations around the background reference and do not represent negative NM content. These values were retained without truncation for statistical analysis to preserve the distribution resulting from background correction. The percentage of cytosolic area occupied by NM was calculated as the percentage of intracellular NM area relative to the total cytosolic area.

For extraneuronal NM quantification, serial H&E-stained, 5 µm-thick sections of the rat SN were collected, with one section every 34^th^, resulting in 5–6 sections per animal encompassing the entire SN, corresponding to half of the full SN series. Extraneuronal NM was identified by the presence of compacted brown NM granules/debris outside neurons. The stained sections were scanned using the Olympus Slideview VS200 slide scanner and visualised with Olyvia 3.3 software. Quantifications were performed using QuPath software, where extraneuronal NM granules were manually identified and counted.

For the quantification of cytoplasmic and nuclear inclusions, four representative 5 µm-thick rat brain sections covering the entire SNpc were selected for immunofluorescence staining with anti-p62 and anti-TH antibodies. Confocal images were captured using the LSM 980 with Airyscan 2, and quantifications were carried out using ZEN 3.1 software. Both p62-positive cytoplasmic and nuclear inclusions were quantified, and their presence within NM/TH-positive or negative neurons was recorded.

For the quantification of neuroinflammation, 5 µm-thick rat paraffin-embedded sections of the SN stained for microglial/macrophage glycoprotein CD68 were scanned using the Olympus Slideview VS200 slide scanner and the Olyvia 3.3 software, with one section containing SN at the same anatomical plane selected per animal. An AI-assisted algorithm in the Olympus VS200 Desktop 3.3 software was implemented for quantifying CD68-positive cells. Data was normalized to the total SN area to minimize bias. Briefly, the classifier was trained to identify CD68-positive profiles within the SN using 14 representative images from different animals spanning the three experimental groups and the observed range of immunoreactivity. Manual annotations established the classification criteria, and the final output was expressed as the number of CD68-positive profiles normalized to SN area. All images included in the analysis were manually reviewed, and classification errors were corrected before final quantification, providing 100% post-classification quality control.

For quantifications performed on a single section, one coronal section at a comparable anatomical level was selected for each rat based on cytoarchitectonic landmarks of the SN. Specifically, we selected the level at which the SN was clearly developed and identifiable, using the surrounding anatomical structures and the organization of the nigral region as landmarks. This level corresponded approximately to AP −6.04 mm relative to Bregma, according to the rat brain atlas of Paxinos and Watson. The same anatomical criterion was applied across all experimental groups, independently of the observed pathological phenotype.

The integrity of STR DA fibers and levels of αSYN were evaluated by quantifying striatal TH or αSYN immunosignal, respectively, by OD. Serial coronal sections covering the whole striatum (10 sections/animal) were selected. TH/αSYN-immunostained 5 µm-thick paraffin-embedded sections were scanned with an Olympus Slideview VS200 slide scanner and visualised using QuPath software. Quantification was performed using Image J. Striatal densitometry values were corrected for non-specific background staining by subtracting densitometric values obtained from the cortex within the same images. Data are expressed as OD or absorbance, calculated based on the logarithmic intensity of light transmission through the material using the formula: 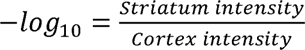.

For image acquisition, immunohistochemical preparations were scanned at 20× magnification, whereas immunofluorescence preparations, including the confocal analyses, were acquired at 40× magnification. For NM analyses, H&E-stained sections were scanned at 20× magnification.

### Chromatographic determination of dopamine metabolites

For Ultra-Performance Liquid Chromatography–Tandem Mass Spectrometry (UPLC-MS/MS) analyses, brain samples were obtained from 24 Sprague-Dawley male rats that were stereotaxically injected in the right SNpc with either AAV-αSYN/vehicle, AAV-TYR/vehicle or AAV-αSYN/AAV-TYR, following above-mentioned procedures. Contra- and ipsilateral ventral midbrain (vMB) were quickly dissected out, frozen on dry ice and stored at -80 °C until analysed.

UPLC-MS/MS analysis of vMB rat samples was performed using our previously validated method (35) with modifications. A stock solution of the internal standard (IS), dopamine-1,1,2,2-d4 hydrochloride (DA-d4) (Sigma-Aldrich, #73483), was prepared in 25 mM formic acid (FA) and stored at -80 °C until use. DA and L-DOPA solutions were freshly prepared, mixed and serially diluted with 25 mM FA to create a concentration series for calibration curves. The IS was added to the mixture at a final concentration of 500 nM. To generate calibration curves and quality control samples, one side of the vMB from three animals was pooled and homogenized in 4 ml of 250 mM FA. Homogenates were divided into 120 µl aliquots, followed by the addition of 40 µM of the mixture. Samples were then centrifuged at 20,000 x g for 10 min at 4 °C, and 120 μl of the supernatant combined with an equal volume of 25 mM FA, was transferred to an Ostro^®^ protein precipitation and phospholipid removal plate (Waters, #186005518) and filtered. Finally, 7 μl were injected into the UPLC-MS/MS system. For brain samples, calibration levels spanned a linear range of 1–3000 nM for each metabolite (DA and L-DOPA), maintaining the IS concentration at 500 nM. On the day of analysis, frozen brain samples were homogenized in 300 µl of 250 mM FA. A 10 µl aliquot was taken and diluted 1:10 with FA for protein quantification using the BCA method. IS was then added to samples at a final concentration of 500 nM. Samples were centrifuged at 20,000 x g for 10 min at 4 °C, and the resulting supernatant was filtered using the Ostro^®^. Finally, 7 μl of the processed sample was injected into the UPLC-MS/MS system for analysis.

An Acquity UPLC system coupled to a Xevo TQ-S triple quadrupole mass spectrometer with an electrospray ionization interface was utilized. Instrument control, data acquisition, and analysis were conducted using MassLynx v4.1 software. Chromatographic separation was achieved using an Acquity™ HSS T3 column (1.8 μm, 2.1 × 100 mm) (Waters, #186003539) paired with an Acquity™ HSS T3 VanGuard pre-column (100 Å, 1.8 μm, 2.1 × 5 mm) (Waters, #186003976). The column was maintained at 40 °C, while the autosampler was held at 6 °C. The mobile phase consisted of solvent A (methanol 100%) and solvent B (25 mM Fa in water), delivered at a flow rate of 0.4 ml/min with the following gradient profile: 0.5% B maintained for 0.5 min, 8% B at 2.6 min, 55% B at 2.9 min, 60% B at 3.3 min, 80% B at 4.3 min, 90% B at 4.4 min and maintained for 0.5 min, and 0.5% B at 5 min followed by 1 min of equilibration. The mass spectrometer detector operated under the following parameters: source temperature 150 °C, desolvation temperature 450 °C, cone gas flow 50 l/h, desolvation gas flow 1100 l/h and collision gas flow 0.15 ml/min. Argon was employed as the collision gas. The capillary voltage was set at 0.5 kV. The electrospray ionization source was alternated between positive and negative ion modes, depending on the analyte. Multiple Reaction Monitoring (MRM) acquisition settings for the targeted metabolites are summarized in **Error! Reference source not found.**4. Samples were considered acceptable if their concentrations were between the limit of detection (LOD) and the limit of quantification (LOQ) or exceeded the LOQ. Concentrations below the LOD were reported as the LOD value. DA vesicular uptake was calculated as DA:L-DOPA ratio (36). Because DA and L-DOPA were quantified in vMB samples, the resulting DA:L-DOPA ratio represents the sampled vMB as a whole and does not provide region-resolved measurements of SNpc versus VTA dopamine handling.

**Table 4.** MRM acquisition settings.

| Analyte | Transition (m/z) | Retention time (min) | Dwell time (ms) | Cone voltage (V) | Collision energy (eV) | Capillary voltage (kV) |
|---|---|---|---|---|---|---|
| DA-4d (IS) | 157.83>9.8 | 1.44 | 52 | 10 | 20 | 0.5 |
| DA | 153.93 > 90.57 | 1.46 | 52 | 10 | 20 | 0.5 |
| L-DOPA | 198,1 > 152.1 | 1.48 | 52 | 15 | 15 | 0.5 |

### Statistical analyses

For the studies in rodents, statistical analyses were performed using GraphPad Prism (version 8, GraphPad Software Inc., USA). No statistical methods were used to pre-determine sample size but sample sizes are equivalent to those reported in previous similar publications (12,24,25). Data are presented as histogram bars, with all values expressed as the mean ± standard error of the mean (SEM). Outlier values were identified using the ROUT method, with a false discovery rate threshold of Q = 1%, and excluded from the analyses when applicable. Normality of data distribution was assessed using the Shapiro-Wilk test. For comparisons between two groups, an unpaired t-test was used for normally distributed data, while the Mann-Whitney test was applied for non-parametric data. For comparisons among three groups or more involving a single variable, a one-way ANOVA followed by Holm-Sidak’s post-hoc test was performed for normal data, or a Kruskal-Wallis test with Dunn’s post-hoc for non-parametric data. For comparisons involving two variables, a two-way ANOVA followed by Tukey’s post-hoc test was applied. For comparisons involving three variables, a three-way ANOVA followed by Tukey’s post-hoc test was used. In all analyses, the null hypothesis was rejected at the 0.05 level.

For the studies in humans, to assess differences in the quantity of human dopamine neurons in different regions and between controls and PD cases, multivariate linear regression tests were applied covarying for age, sex, and post-mortem delay using IBM SPSS (SPSS Inc., Chicago, IL, USA, Version 29.0.2.0), and *p* < 0.05 was considered statistically significant. Graphs were made with GraphPad Prism10 (GraphPad Prism In., USA, version 10.6.0 (796)). Differential gene expression analyses in TH immunoreactive neurons located in the ventral tier of the SNpc versus the dorsal tier of the SNpc and VTA were performed in the R statistical program using the limma-voom framework (37), controlling for post-mortem delay, RNA integrity (DV200), sex, processing plate, and subject.

In all cases, researchers blinded to the case group conducted data collections.

## Results

### Increased αSYN (*SNpcA*) mRNA expression in vulnerable human substantia nigra region enriched for NM-laden neurons

We first assessed a potential link between αSYN expression levels and the vulnerability of NM-laden neurons in control human midbrains, separating the data by regional susceptibility to degeneration in PD: the vulnerable SNv versus the more resistant SNd and VTA (Fig. 1a,b). To this end, we analysed mRNA expression levels of *SNpcA* and *TH* in DA neurons from these regions. There was slightly less *TH* mRNA expression in the SNv compared with the SNd/VTA (Fig. 1f) but much greater *SNpcA* mRNA expression (two-fold higher in DA neurons of the SNv compared with the SNd/VTA, Fig. 1g). While more *SNpcA* than *TH* mRNA was observed in control SNv DA neurons, there was a close correlation between the expression levels of *TH* and *SNpcA* in these SNv neurons (Fig. 1h). This correlation was not observed in SNd/VTA DA neurons with a high proportion of neurons expressing higher levels of *TH* compared with *SNpcA* mRNA (Fig. 1h). This data identifies significant differences between the regional expression levels of *TH* and *SNpcA* mRNAs in midbrain DA neurons in healthy human controls, with the vulnerable SNv having higher *SNpcA* and slightly lower *TH* mRNA expression compared with the more resistant SNd/VTA DA neurons.

**Figure 1.**
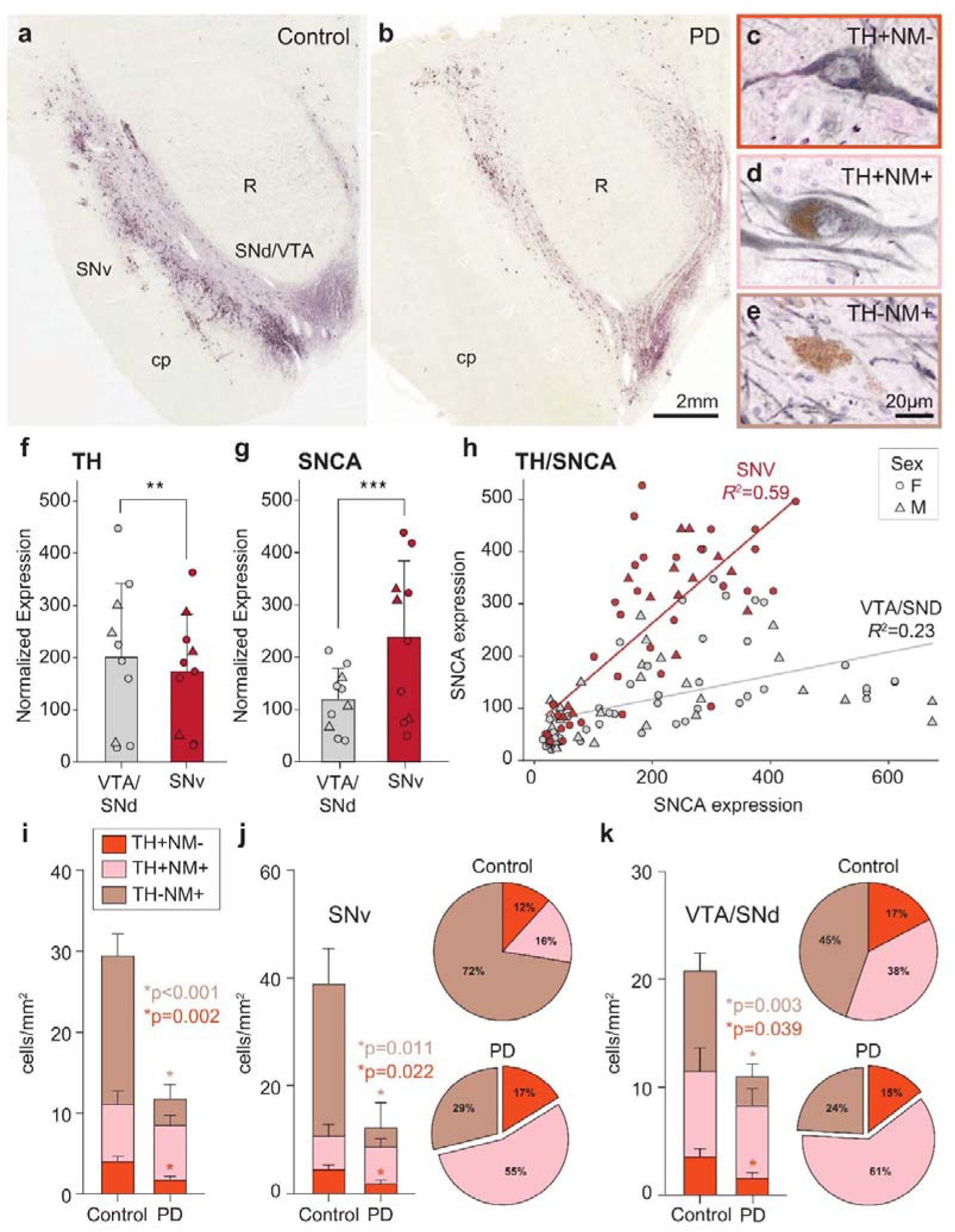
Increased SNpcA mRNA expression in vulnerable human substantia nigra region enriched for NM-laden neurons. **(a, b)** Representative low magnification photomicrographs of the DA cell regions (TH-immunolabelled) in the human midbrain (medial at right, ventral at bottom) in a control (a) and PD patient (b) showing the relative selective loss of the ventral tier of the SNpc (SNv). cp=cerebral peduncle, R=red nucleus, SNd/VTA=dorsal tier of the SNpc and ventral tegmental area. **(c-e)** Representative high magnification photomicrographs of the different types of DA neurons in the human midbrain in TH immunolabelled sections: TH^+^ only neuron (c), NM^+^TH^+^ neuron (d, NM=brown intracytoplasmic NM granules), and NM^+^ only neuron (e). **(f, g)** Quantification of the average TH (f) and SNpcA (g) mRNA expression within the SNv and SNd/VTA in control subjects showing lower TH and higher SNpcA in the SNv; n=10, mean ± standard deviation shown, limma voom, **False Discovery Rate (FDR) adjusted p-value= 0.009, ***FDR adjusted p-value= 4.1 × 10^-9^. **(h)** Correlations between TH and SNpcA mRNA expression in SNv versus SNd/VTA, showing a stronger correlation between transcripts in the SNv (n=10, ROIs = 49, p-value = 1.12 × 10^-10^) than the SNd/VTA (n=10, ROIs = 80, p-value = 6.97× 10^-6^). **(i-k)** Density and proportion of the different DA cell types in controls and PD cases (i) and within each region (j,k); control n=7, PD n=13, mean ± standard deviation shown, multivariate linear regression test was applied to compare cell loss covarying for age, sex, and post-mortem delay. There was a significant loss of DA cells in PD compared to controls (i, F_group_ (df4)=7.079, p<0.001, F_DA cell density_ (df1)=22.737, p<0.001) but no loss of TH^+^NM^+^ neurons (i, F_TH+NM+_ (df1)=0.017, p=0.895). Within the subregions, the most consistent loss was of NM+ only DA neurons in PD (j,k, F_group*subregion_ (df4)=2.058, p=0.1, F_NM+ only_ (df1)=7.882, p=0.007).

We next quantified the total number of DA neurons in these regions of the human midbrain sections analysing three types of DA cells: NM^+^ only, TH^+^NM^+^ and TH^+^ only (Fig. 1c-e). Assessing the regional numbers and types of DA neurons in control sections identified that the SNv had higher DA neuronal densities compared to SNd/VTA (average 39 versus 21 DA cells/mm^2^, F_subregion_ (df4)=5.05, *p*=0.04, F_DA cell density_ (df1)=8.03, *p*=0.02) and a trend for more NM^+^ only DA neurons (average 72% versus 45% NM^+^ only of DA neurons, F_subregion_ (df3)=4.761, *p*=0.04, F_NM only_ (df1)=4.44, *p*=0.06). This data confirms that high NM^+^ DA neurons with low TH immunoreactivity are a feature of the more selectively vulnerable SNv DA cell region in healthy controls (19,38).

Lastly, we analysed which cell types were most affected in the different regions in PD compared to controls. As expected, there was a significantly greater loss of DA neurons in the SNv compared with the SNd/VTA in the PD cases (average 70% versus 40% DA cell loss from controls, F_group*subregion_ (df3)=4.356, *p*=0.01, F_DA cell density_ (df1)=11.93, *p*=0.002). The DA cell type most affected was those NM^+^ only, with no loss of DA neurons containing both NM and TH (Fig. 1i). The loss of NM^+^ only DA neurons in PD was observed in both the SNv and SNd/VTA regions but, as this cell type dominated the SNv, the effect of PD concentrated in this DA cell region (Fig. 1a,b,j,k). DA neurons expressing only TH and no NM were also reduced in PD, but this DA cell type was a small proportion of the total population of DA neurons in the midbrain (<20%, Fig. 1i-k). Our data confirm those of others (19,38) reporting a concentration of NM^+^ DA neurons in the SNv that are more selectively vulnerable to PD compared to DA neurons containing both NM and TH. Overall, our results in humans suggest that increased regional *SNCA* expression aligns with enrichment of NM-laden neurons in the SNv, the region more susceptible to degeneration in PD.

### αSYN overexpression enhances NM-linked neurodegeneration in TYR-expressing rats

To experimentally assess the effects of increased αSYN expression on NM-laden neurons *in vivo*, we combined αSYN overexpression with TYR-induced NM production in rats. Adult male rats received single unilateral stereotaxic injections of either an AAV expressing human αSYN (AAV-αSYN), human TYR (AAV-TYR), or a combination of both (AAV-αSYN + AAV-TYR) above the right SNpc (Fig. 2). Rats were euthanized at 2, 4, and 15 months (m) post-injection and processed for histological characterization (Fig. 2a). Ipsilateral nigrostriatal expression of human αSYN was confirmed by immunohistochemistry using a human-specific αSYN antibody in both αSYN-injected and αSYN+TYR co-injected animals (Fig. 2b). αSYN co-expression with TYR did not alter TYR expression (Fig. 2c). No expression of human αSYN or TYR was detected in the contralateral, non-injected hemispheres of any experimental group.

**Figure 2.**
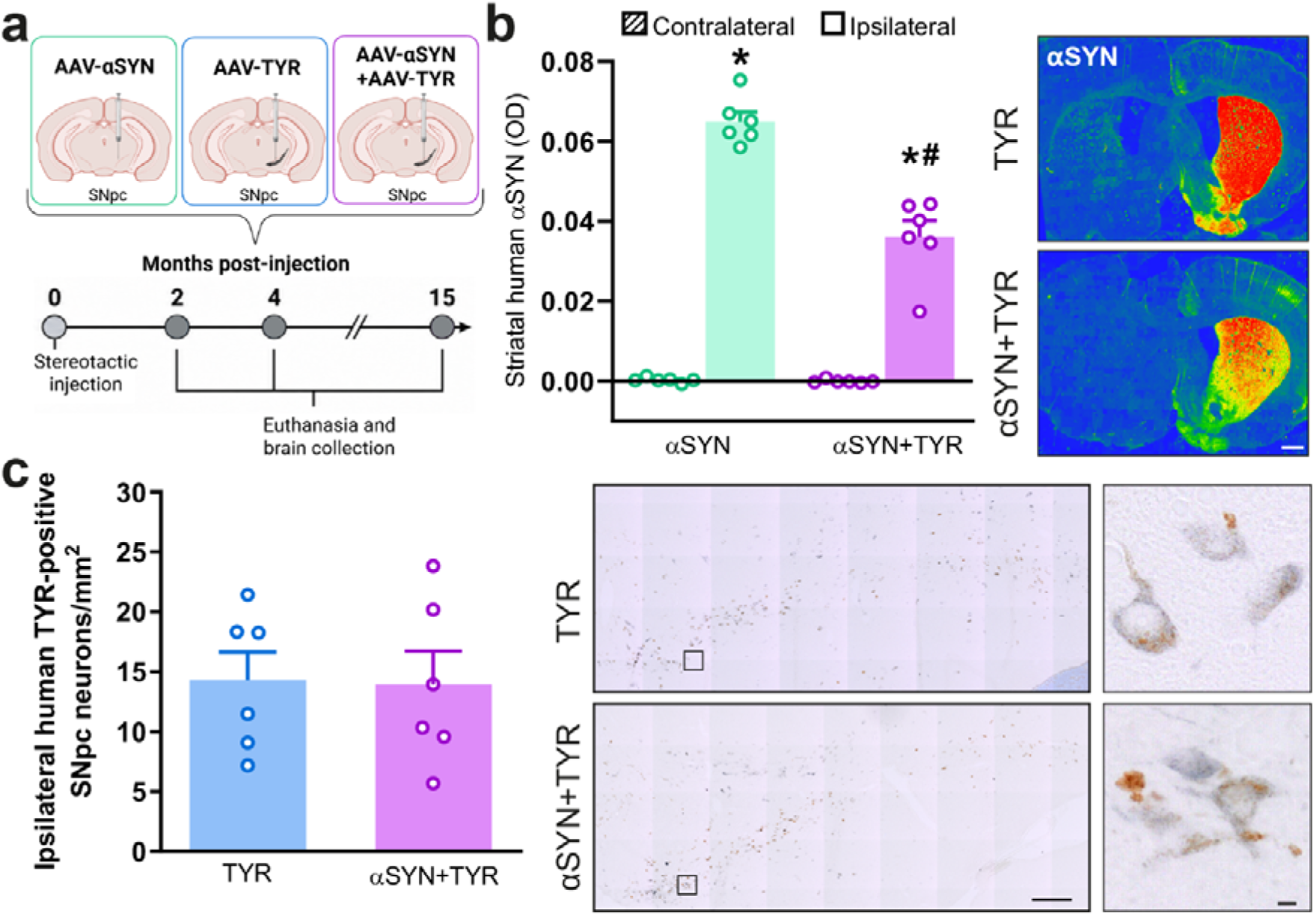
Overexpression of αSYN and/or TYR in the nigrostriatal pathway of rats. (**a**) Schematic representation of the experimental design. (**b**) Left: Optical density (OD) of striatal αSYN in αSYN- and αSYN+TYR-expressing rats at 2 months (m) post-AAV injection. Two-way ANOVA with Tukey’s post hoc test: treatment, F(1,20) = 38.1; hemisphere, F(1,20) = 453.3; treatment × hemisphere, F(1,20) = 36.6; all p < 0.0001. *p<0.05 compared to the respective contralateral hemisphere; ^#^p<0.05 compared to ipsilateral αSYN-expressing animals. Right: Representative photomicrographs of striatal αSYN immunolabeling. Scale bar, 1 mm. (**c**) Left: Quantification of ipsilateral TYR-positive SNpc neurons in TYR- and αSYN+TYR-expressing rats at 2 m post-AAV injection (Mann-Whitney test). Right: Representative photomicrographs of ipsilateral SNpc TYR immunolabeling (TYR-positive neurons, blue; unstained NM, brown). Scale bars, 200 µm and 5 µm (inset). The schematic in (**a**) was created with BioRender.com. In (b) and (c), n=6 animals per experimental group.

We have previously reported that experimentally inducing NM production in TYR-expressing rodents is ultimately associated with nigrostriatal degeneration (12,23–25). Here, we examined the impact of αSYN overexpression on the extent and temporal progression of NM-linked neurodegeneration. First, striatal TH-positive fibers were quantified over time by optical densitometry (OD) across all experimental animal groups (Fig. 3a). In αSYN-expressing animals, which lack NM, the ratio of ipsilateral to contralateral striatal TH□O□D remained relatively stable over time, with no significant fiber loss (Fig. 3a). As previously reported (12), NM-producing TYR-expressing animals showed a significant reduction in ipsilateral striatal TH fibers starting at 4 m post-AAV injection, corresponding to ∼61% fiber loss relative to the contralateral side (Fig. 3a). This degeneration was further aggravated in αSYN+TYR-expressing rats, which combine NM production with increased αSYN levels, reaching up to ∼85% fiber loss by 15 m (Fig. 3a). Indeed, the temporal evolution of striatal denervation differed between the TYR and αSYN+TYR groups. Although αSYN+TYR rats showed less TH-fiber loss than TYR rats at 4 months, this difference was transient, as TH-fiber loss subsequently progressed in αSYN+TYR animals to reach the lowest mean striatal TH signal at 15 months, whereas the loss in TYR animals remained relatively stable. To determine whether the enhanced reduction of nigrostriatal fibers observed in αSYN+TYR-expressing animals at 15 m was associated with increased loss of SNpc neurons, we quantified TH-positive nigral cell bodies at this particular time-point in all animal groups (Fig. 3b). Both TYR- and αSYN+TYR-expressing rats exhibited a significant reduction in TH-positive neurons in the ipsilateral SNpc, compared either to their respective contralateral SNpc or to the ipsilateral SNpc of αSYN-only expressing animals (Fig. 3b). As observed in humans (Fig. 1), part of this reduction was attributable to a phenotypic downregulation of TH protein expression within NM-laden neurons (Fig. 3c). To distinguish between TH downregulation and actual neuronal loss, we quantified the total number of SNpc DA neurons, including TH^+^NM^−^, TH^+^NM^+^, and TH^−^NM^+^ cells, which revealed a significant ipsilateral loss of total SNpc DA neurons in αSYN+TYR-expressing rats compared to either αSYN- or TYR-expressing groups alone (Fig. 3d). Similar to humans (Fig. 1), neurons with downregulated TH expression (TH^−^NM^+^) were the most vulnerable to neurodegeneration in αSYN+TYR-expressing rats (Fig. 3e). Supporting the occurrence of enhanced neurodegeneration in αSYN-overexpressing NM-laden neurons, extraneuronal NM debris, released from dying neurons, was increased in αSYN+TYR-expressing animals compared to TYR-expressing rats (Fig. 3f). This was associated with heightened macrophage activation, as evidenced by increased CD68 immunoreactivity surrounding NM debris (Fig. 3g). Overall, these results indicate that αSYN overexpression aggravates the degeneration of NM-laden nigrostriatal neurons, but does not affect these neurons in the absence of NM.

**Figure 3.**
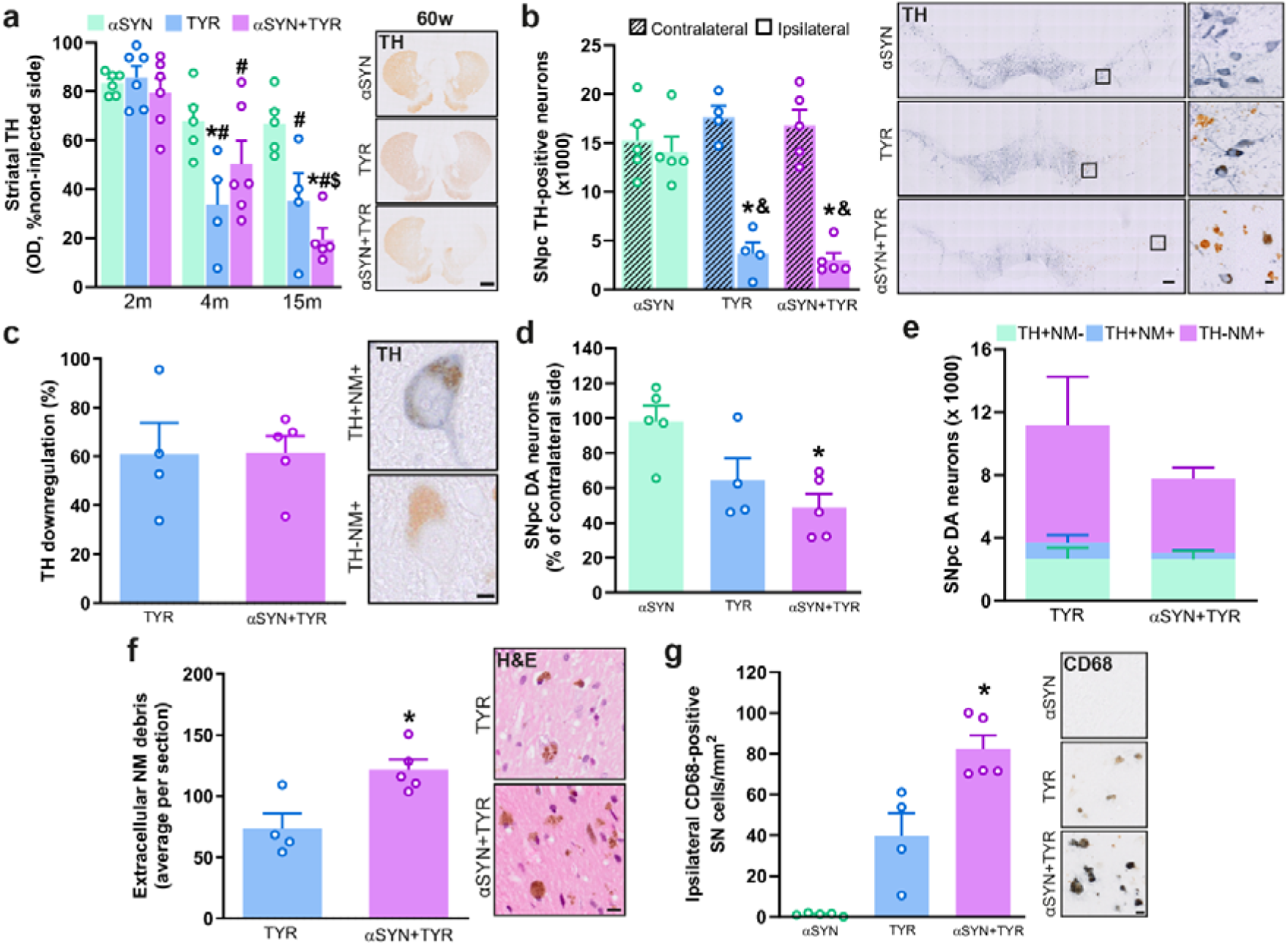
αSYN overexpression enhances nigrostriatal degeneration in NM-producing TYR-expressing rats. (**a**) Left, OD quantification of striatal TH fibers in αSYN-, TYR- and αSYN+TYR-expressing rats at 2, 4 & 15 m post-AAV injection. Right, representative photomicrographs of striatal TH immunolabeling at 15 m. Scale bar, 1 mm. (**b**) Left, Quantification of SNpc TH-positive neurons in αSYN-, TYR- and αSYN+TYR-expressing rats at 15 m. Right, Representative photomicrographs of TH immunolabeling in SNpc. Scale bars, 200 µm & 10 µm (inset). (**c**) Left, TH^−^NM^+^ neurons versus total NM^+^ neurons (including TH^+^NM^+^ and TH^−^NM^+^ neurons) as an index of TH downregulation in TYR- and αSYN+TYR-expressing rats at 15 m. (**d**) Quantification of total SNpc DA neurons in αSYN-, TYR- and αSYN+TYR-expressing rats at 15 m. (**e**) Density and proportion of the different DA cell types in TYR- and αSYN+TYR-expressing rats at 15 m. (**f**) Left, quantification of extraneuronal NM debris in TYR- and αSYN+TYR-expressing rats at 15 m. Right, representative photomicrographs of ipsilateral extraneuronal NM debris (arrowheads) in H&E-stained sections. Scale bar, 10 µm. (**g**) Left, quantification of ipsilateral CD68-positive SNpc cells in αSYN-, TYR- and αSYN+TYR-expressing rats at 15 m. Right, representative photomicrographs of CD68 immunolabeling in SNpc. Scale bar, 10 µm. In (**b, c** & **g)**, immunolabeling is blue and unstained NM is brown. n = 4–6 (**a**) or 4–5 (**b–g**) animals per group. Two-way ANOVA with Tukey’s post hoc test (**a, b**): (**a**) treatment, F(2,38) = 10.7, p = 0.0002; time, F(2,38) = 33.1, p < 0.0001; treatment × time, F(4,38) = 4.3, p = 0.0055; (**b**) treatment, F(2,22) = 7.1, p = 0.0041; hemisphere, F(1,22) = 71.4, p < 0.0001; treatment × hemisphere, F(2,22) = 14.4, p < 0.0001. Mann–Whitney test (**c, f**); Kruskal–Wallis test with Dunn’s post hoc test (**d, g**). In a, b, d, g, *p < 0.05 vs respective αSYN; ^#^p < 0.05 vs respective 2 m; ^$^p < 0.05 vs respective 4 m; ^&^p < 0.05 versus respective contralateral side. In f, *p < 0.05 vs TYR.

### αSYN overexpression enhances NM-linked inclusion formation in TYR-expressing rats

We have previously reported that progressive NM accumulation in TYR-expressing rodents is associated with the formation of intracellular inclusion bodies, similar to those found in PD and aging human brains (12,23,24). These include, respectively, cytoplasmic LB-like inclusions and nuclear Marinesco bodies (MB), both of which are immunopositive for p62, a common component of neuropathological inclusions and an indicator of general proteostasis failure (15). We therefore examined whether the enhanced degeneration of NM-laden neurons induced by αSYN overexpression is associated with greater inclusion formation in these cells. As previously described (12), AAV-TYR-injected rats exhibited p62-immunopositive cytoplasmic inclusions in nigral NM-laden neurons, which peaked at 2 m post-AAV-TYR injection and subsequently decreased from 4 m onward, in parallel with neurodegeneration (Fig. 4a). In contrast, αSYN+TYR-co-expressing rats showed a more sustained accumulation of p62-positive cytoplasmic inclusions, which extended up to 4 m before their levels were decreased by 15 m post-AAV injection (Fig. 4a). Regarding MB-like inclusions, a progressive accumulation of intranuclear p62-positive inclusions was observed mainly in αSYN+TYR-co-injected animals, starting from 4 m onward (Fig. 4b). Notably, in all experimental groups, p62-positive inclusions, both cytoplasmic and nuclear, were restricted to melanized neurons and were not observed in αSYN-only expressing animals lacking NM formation.

**Figure 4.**
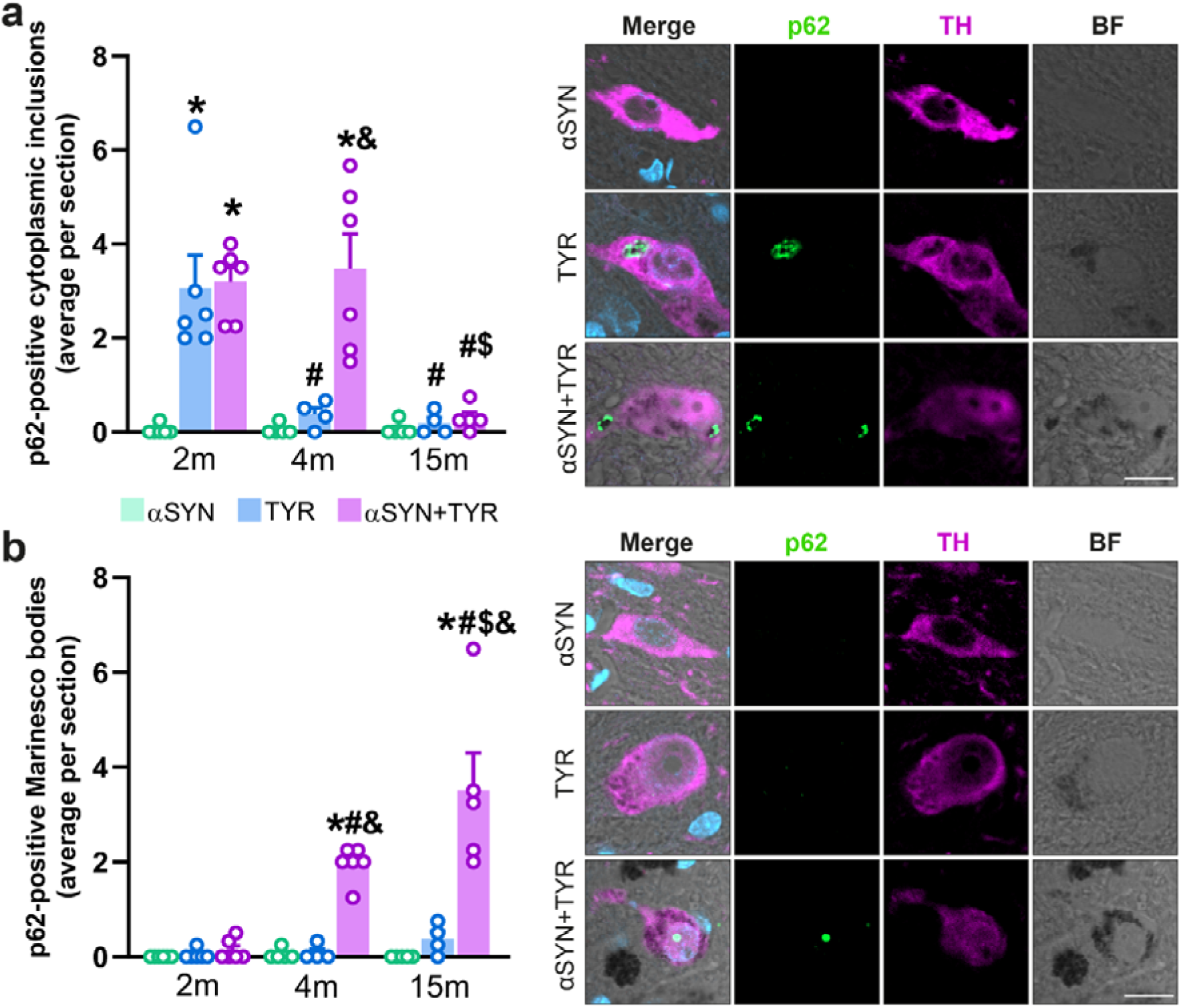
αSYN overexpression enhances NM-linked inclusion formation in TYR-expressing rats. (**a, b**) Left, quantification of p62-positive cytoplasmic (a) and nuclear (b) SNpc inclusions in αSYN-, TYR- and αSYN+TYR-expressing rats at 2, 4 & 15 m post-AAV injection. Right, representative photomicrographs of ipsilateral p62-positive cytoplasmic inclusions. p62, green; TH, magenta; DAPI, blue; NM, dark grey in brightfield (BF). Scale bar, 10 µm. n=4-6 animals per group. Two-way ANOVA with Tukey’s post hoc test: (**a**) treatment, F(2,38) = 23.18; time, F(2,38) = 15.7; treatment × time, F(4,38) = 8.3; (**b**) treatment, F(2,38) = 45.4; time, F(2,38) = 15.9; treatment × time, F(4,38) = 12.5; all p < 0.0001. *p < 0.05 vs respective αSYN; #p < 0.05 vs respective 2 m; $p < 0.05 vs respective 4 m; &p < 0.05 vs respective TYR.

Next, we assessed intracellular αSYN accumulation by counting total human-specific αSYN-positive SNpc neurons in the ipsilateral side of αSYN- and αSYN+TYR-injected rats at 2, 4 and 15 m post-AAV injection (Fig. 5a). αSYN+TYR co-injected animals displayed fewer αSYN-positive SNpc neurons than αSYN-injected animals at all time-points examined (Fig. 5a). Phosphorylation of αSyn (pαSyn) at serine 129 (Ser129) is widely considered a hallmark of pathological αSyn in PD and related synucleinopathies (39,40). We thus quantified ipsilateral pαSyn-positive SNpc neurons over time across all experimental groups (Fig. 5b).

**Figure 5.**
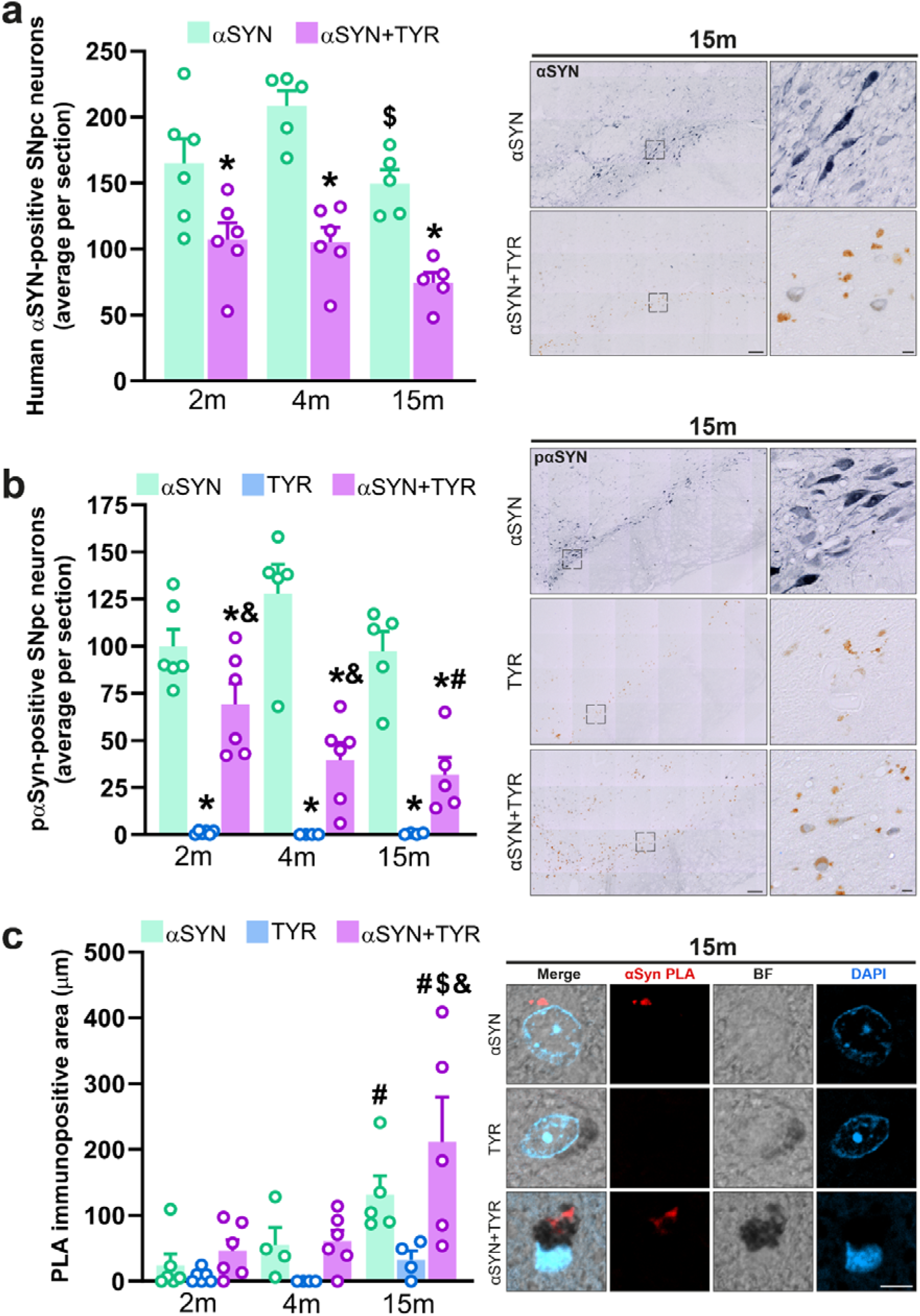
αSYN overexpression enhances αSyn oligomerization in NM-producing rats. (**a– c**) Left, quantification of human αSYN-positive neurons (**a**), pαSYN-positive neurons (**b**) and oligomeric αSYN PLA-immunopositive area (**c**) in the SNpc of αSYN-, TYR- and αSYN+TYR-expressing rats at 2, 4 & 15 m post-AAV injection (αSYN and αSYN+TYR only in **a**). Right, representative photomicrographs of ipsilateral human αSYN immunolabeling (**a**), pαSYN immunolabeling (**b**) and αSyn PLA signal at 15 m. Scale bars, 10 µm (a,b) and 5 µm (c). n=4-6 animals per group. Two-way ANOVA with Tukey’s post hoc test: (**a**) treatment, F(1,27) = 54, p < 0.0001; time, F(2,27) = 5.6, p = 0.0093; treatment × time, F(2,27) = 1.6, p = 0.2279; (**b**) treatment, F(2,38) = 93.6, p < 0.0001; time, F(2,38) = 1.8, p = 0.1753; treatment × time, F(4,38) = 3, p = 0.0293; (**c**) treatment, F(2,37) = 8.2, p = 0.0011; time, F(2,37) = 10.8, p = 0.0002; treatment × time, F(4,37) = 1.8, p = 0.1466. *p < 0.05 vs respective αSYN; #p < 0.05 vs respective 2 m; $p < 0.05 vs respective 4 m; &p < 0.05 vs respective TYR.

Using an antibody that detects both human and murine pαSyn, only a few SNpc neurons were immunopositive in TYR-expressing animals at any time point (Fig. 5b). In contrast, all αSYN-expressing animals, alone or in combination with TYR, exhibited pαSyn-positive nigral neurons with diffuse cytosolic pαSyn immunoreactivity (Fig. 5b). In αSYN+TYR-expressing rats, the number of pαSyn-positive neurons declined over time, starting at 2 m post-injection (Fig. 5b). Given that αSyn oligomers are considered toxic intermediates in PD-related neurodegeneration (41,42), we quantified αSyn oligomeric species in these animals using the proximity ligation assay (PLA), a method previously validated for detecting αSyn oligomers in post-mortem PD brains (43). In parallel with the decrease in total αSYN and pαSyn immunostaining, αSYN+TYR-expressing rats exhibited a corresponding increase in αSyn oligomeric species, detectable up to 15 m post-AAV injection (Fig. 5c).

Overall, these results indicate continuous αSYN oligomerization and sustained p62-positive inclusion formation in NM-laden neurons overexpressing αSYN. Furthermore, the reductions in total αSYN- and pαSyn-positive SNpc neurons in αSYN+TYR-expressing rats suggest that NM-laden neurons accumulating total αSYN and pαSyn may be the ones most vulnerable to degeneration.

### αSYN overexpression enhances NM accumulation in TYR-expressing rats

While experimentally inducing NM production in TYR-expressing rodents is sufficient to stimulate αSYN pathology and neurodegeneration (12,24,25), it is unknown whether αSYN overexpression influences NM accumulation, which could contribute to the enhanced NM-associated pathology observed in αSYN+TYR-expressing animals. To examine the potential effects of αSYN upregulation on NM accumulation, we assessed intracellular NM levels in the SNpc of αSYN+TYR-expressing rats compared with TYR-only expressing animals, since αSYN expression alone does not produce NM in the absence of TYR. Intracellular NM density and the percentage of NM-occupied cytosolic area were quantified in H&E-stained nigral sections from TYR- and αSYN+TYR-expressing animals at 2, 4, and 15 m post-AAV injections. Compared to TYR-only expressing animals, αSYN+TYR-expressing rats exhibited a significant increase in both intracellular NM density and the percentage of cytosolic area occupied by NM from 4 m onward (Fig. 6 and Supplementary Fig. 2). These findings indicate that αSYN overexpression accelerates intracellular NM accumulation in SNpc neurons of TYR-expressing rats.

**Figure 6.**
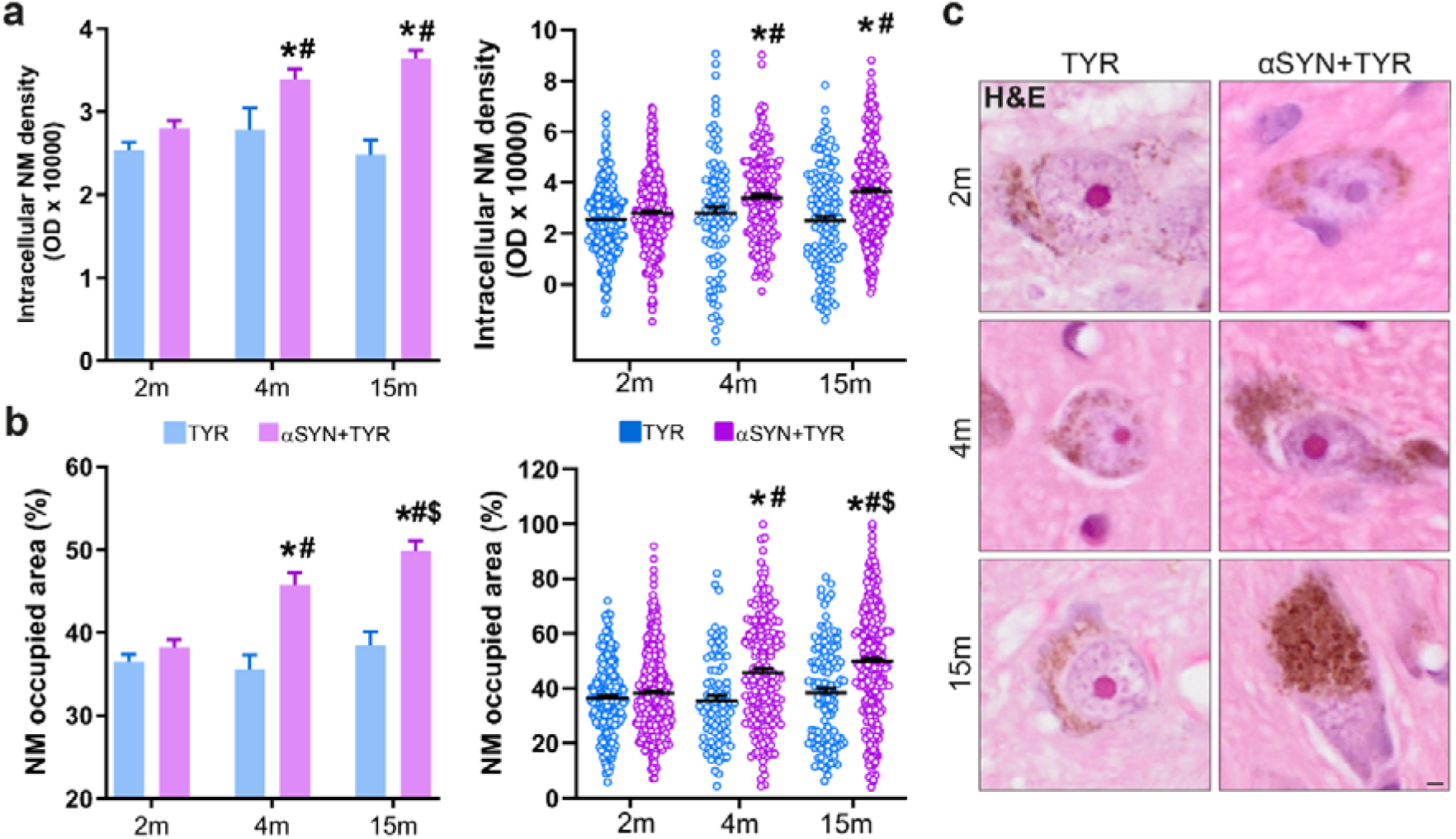
αSYN overexpression enhances NM accumulation in TYR-expressing rats. (**a,b**) Intracellular NM OD (**a**) and percentage of neuronal cytosolic area occupied by NM (**b**) in ipsilateral SNpc DA neurons from TYR- and αSYN+TYR-expressing rats at 2, 4, and 15 m post-AAV injection. Left: mean ± SEM; right: individual values. (**c**) Representative photomicrographs of H&E-stained brain sections from the different experimental groups (unstained NM in brown). Scale bar, 2 µm. Two-way ANOVA with Tukey’s post hoc test (**a, b**): (**a**) treatment, F(1,1203) = 38.4, p < 0.0001; time, F(2,1203) = 7.5, p = 0.0006; treatment × time, F(2,1203) = 6.8, p = 0.0012; (**b**) treatment, F(1,1203) = 49.0, p < 0.0001; time, F(2,1203) = 15.5, p < 0.0001; treatment × time, F(2,1203) = 9.4, p = 0.0001. Sample sizes for **a, b** (neurons/rats at 2, 4 & 15 m, respectively): TYR, 227/6, 89/4 and 118/4; αSYN+TYR, 294/6, 190/5 and 291/5. *p<0.05 vs respective TYR; #p<0.05 vs respective 2 m; $p<0.05 vs respective 4 m.

**Supplementary Figure 2.**
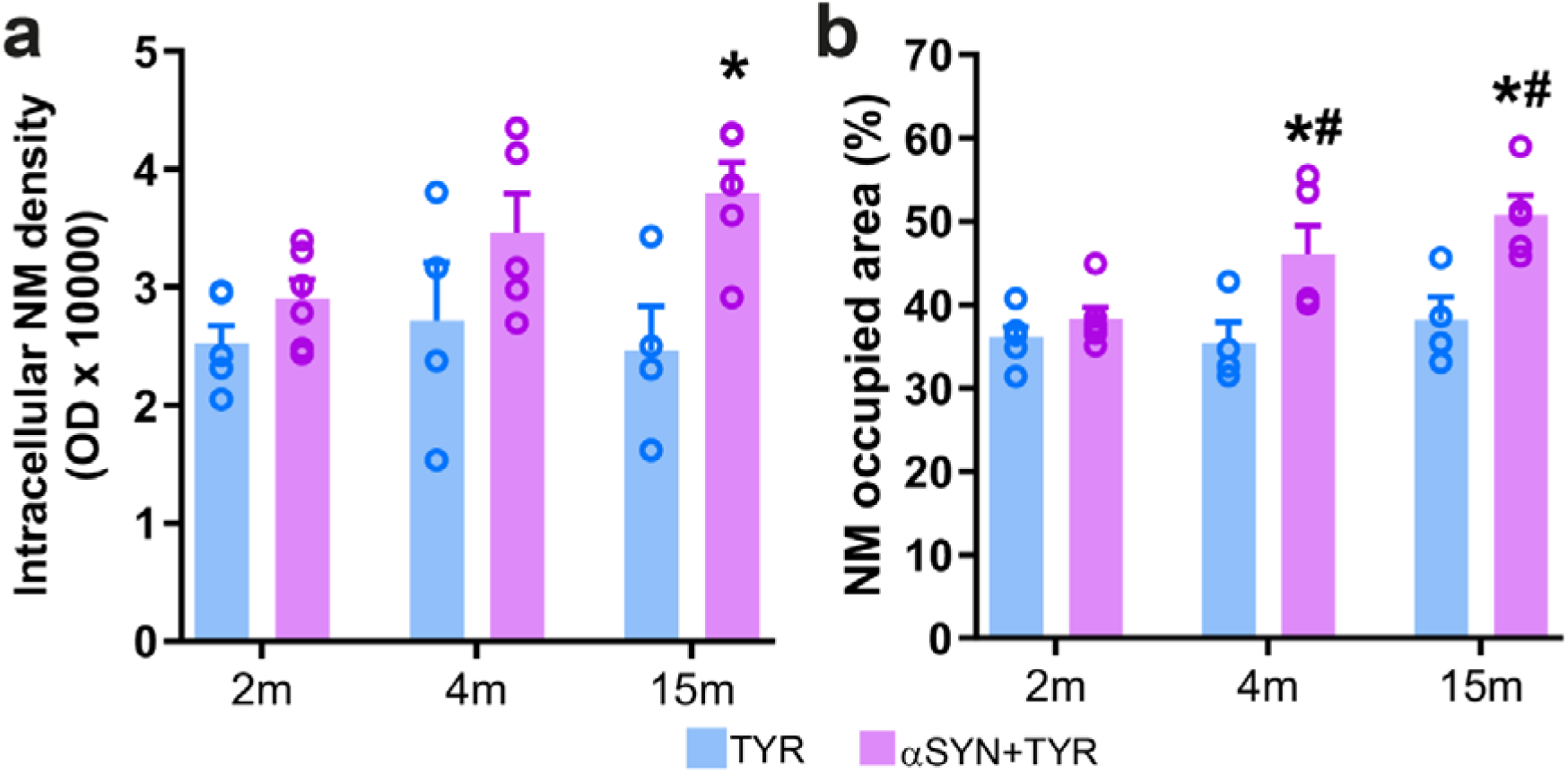
Intracellular NM levels per animal. Intracellular NM OD (**a**) and percentage of neuronal cytosolic area occupied by NM (**b**) in ipsilateral SNpc DA neurons from TYR- and αSYN+TYR-expressing rats at 2, 4 & 15 m post-AAV injection. Two-way ANOVA with Tukey’s post hoc test: (**a**) treatment, F(1,24) = 12.3, p = 0.0018; time, F(2,24) = 1.4, p = 0.2548; treatment × time, F(2,24) = 1.5, p = 0.2524; (**b**) treatment, F(1,24) = 20.2, p = 0.0002; time, F(2,24) = 5.2, p = 0.0132; treatment × time, F(2,24) = 3.2, p = 0.0607. Sample sizes (rats at 2, 4 & 15 m, respectively): TYR, 6, 4 and 4; αSYN+TYR, 6, 5 and 5. *p<0.05 vs respective TYR; #p<0.05 vs respective 2 m.

We next investigated the potential mechanisms by which αSYN overexpression leads to increased intracellular NM levels in TYR-expressing animals. αSYN oligomeric species, which are increased in αSYN+TYR-expressing rats (Fig. 5c), are known to bind to lipids and permeabilize intracellular organelle membranes, including those of mitochondria, lysosomes, and synaptic vesicles (44–46). αSYN-mediated permeabilization of DA-containing vesicles may result in the leakage of stored DA into the cytosol, where it can undergo oxidation to form NM. Supporting this concept, αSYN overexpression increases cytosolic DA levels in chromaffin cells (47), while overexpression of vesicular monoamine transporter 2 (VMAT2) in TYR-expressing rats enhances DA vesicular encapsulation, reducing free cytosolic DA and its subsequent conversion into NM (24). Based on these findings, we hypothesized that αSYN oligomerization in αSYN+TYR-expressing animals may induce the permeabilization of DA-containing vesicles, thereby increasing cytosolic DA available for oxidation into NM. To evaluate the integrity of the DA vesicular system, we first assessed DA vesicular uptake using the ratio of DA to L-DOPA levels measured by UPLC-MS/MS (24) in αSYN-, TYR-, and αSYN+TYR-expressing rats (Fig. 7a). αSYN overexpression alone did not affect DA vesicular uptake, whereas TYR-expressing animals exhibited a significant decrease (Fig. 7b). Notably, co-expression of αSYN and TYR further reduced DA vesicular uptake compared to either αSYN- or TYR-only expressing rats (Fig. 7b).

**Figure 7.**
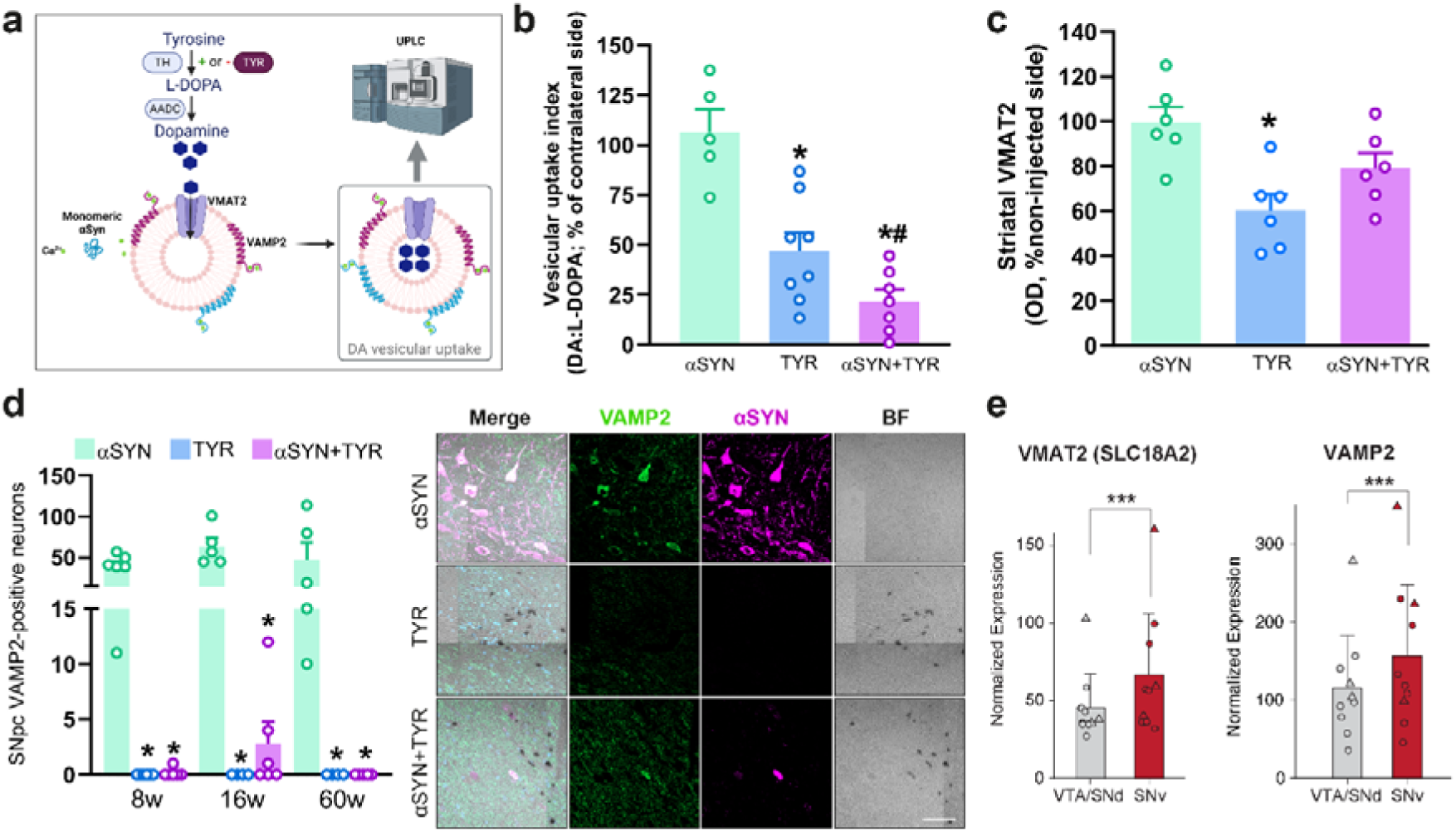
αSYN overexpression enhances NM-linked impairment of vesicular dopamine handling in TYR-expressing rats. **(a)** Schematic of the vesicular DA uptake assay in the rat SN. DA is transported into synaptic vesicles by VMAT2, and vesicular DA uptake index is quantified by UPLC-MS/MS. Created with BioRender.com. **(b)** UPLC-MS/MS quantification of the ipsilateral-to-contralateral DA vesicular uptake index, calculated as the DA:L-DOPA ratio, in αSYN-, TYR-, and αSYN+TYR-expressing rats at 2 months post-AAV injection. **(c)** Striatal VMAT2 immunoreactivity in αSYN-, TYR-, and αSYN+TYR-expressing rats at 2 months post-AAV injection, expressed as percentage of the contralateral non-injected side. **(d)** Left, quantification of ipsilateral SNpc VAMP2-positive neurons in αSYN-, TYR-, and αSYN+TYR-expressing rats at 2, 4, and 15 months post-AAV injection. Right, representative photomicrographs of ipsilateral VAMP2- and αSYN-positive neurons assessed by immunofluorescence (VAMP2, green; αSYN, magenta; DAPI, blue; NM, dark grey in bright field [BF]). Scale bar, 50 µm. **(e)** Quantification of normalized VMAT2 and VAMP2 mRNA expression in the SNd/VTA and SNv of human control subjects, showing higher expression of both transcripts in the SNv; Data are mean ± standard deviation; circles, females; triangles, males. One-way ANOVA with Holm–Sidak’s (**b**) or Tukey’s (**c**) post hoc test. Two-way ANOVA with Tukey’s post hoc test (**d**): treatment, F(2,38) = 39.8, p < 0.0001; time, F(2,38) = 1.0, p = 0.3753; treatment × time, F(4,38) = 0.7, p = 0.5837. In **e**, limma-voom: FDR-adjusted p = 5.8 × 10□□ for VMAT2 and 2.2 × 10□□ for VAMP2. n=5–8 (**b**), 6 (**c**) and 4–6 (**d**) animals per group; n=10 human subjects (**e**). In **b–d**, *p<0.05 vs respective αSYN; in **b**, #p<0.05 vs respective TYR.

Because reduced vesicular DA storage in TYR-expressing NM-producing rats has previously been associated with VMAT2 downregulation (24), we next assessed whether a similar mechanism could account for the greater impairment of DA vesicular uptake observed in αSYN+TYR-expressing rats. Striatal VMAT2 immunoreactivity was therefore quantified in αSYN-, TYR-, and αSYN+TYR-expressing animals (Fig. 7c). TYR expression significantly reduced striatal VMAT2 levels compared with αSYN alone, consistent with our previous findings (24). In contrast, VMAT2 levels in αSYN+TYR animals were intermediate and did not differ significantly from either αSYN- or TYR-expressing animals. Thus, the greater impairment of DA vesicular uptake in αSYN+TYR animals cannot be explained by a further reduction in VMAT2 levels, suggesting the contribution of additional mechanisms affecting vesicular DA handling.

As αSYN facilitates vesicle maturation and fusion with the plasma membrane through its interaction with vesicle-associated membrane protein 2 (VAMP2) (48), VAMP2 immunoreactivity was examined over time in the SNpc of αSYN-, TYR-, and αSYN+TYR-expressing animals (Fig. 7d). Double immunofluorescence for VAMP2 and αSYN revealed a high number of VAMP2-positive neurons in αSYN-expressing animals, many of which also exhibited αSYN immunoreactivity. In contrast, VAMP2-positive neurons were virtually undetectable in TYR-expressing animals and under baseline conditions in the contralateral hemisphere (not shown). Strikingly, VAMP2-positive neurons also remained virtually undetectable in αSYN+TYR-expressing animals despite αSYN overexpression (Fig. 7d). These findings indicate that the prominent VAMP2 signal associated with αSYN overexpression is lost in the NM-producing environment. To determine whether regional differences in vesicular machinery are also present in the human nigra, we examined VMAT2 (SLC18A2) and VAMP2 expression in the human control GeoMx dataset. Both transcripts were significantly higher in the vulnerable SNv than in the SNd/VTA (Fig. 7e), a region enriched in NM-containing dopaminergic neurons and characterized by higher SNCA expression relative to the SNd/VTA. Notably, the regional co-enrichment of SNCA and VAMP2 in the human SNv parallels the high VAMP2 immunoreactivity observed in αSYN-overexpressing rats, consistent with the known functional relationship between αSYN and VAMP2. These observations suggest a regional association between αSYN and vesicular machinery within the dopaminergic neuronal population most vulnerable to degeneration in PD.

Taken together, these results indicate that increased αSYN levels are associated with greater NM-linked impairment of DA vesicular uptake and with changes suggestive of impaired αSYN-linked VAMP2 stabilization. These alterations are consistent with impaired vesicular DA handling that may increase the cytosolic pool of DA available for oxidation into NM, thereby contributing to increased NM accumulation and associated pathology.

## Discussion

This study supports a pathogenic bidirectional relationship between αSYN and NM that may contribute to nigral neurodegeneration in PD. Our data from human brains demonstrate regional αSYN overexpression in vulnerable pigmented neurons of the SNpc, while our in vivo data indicate that increased αSYN levels accelerate NM formation and NM-associated pathology in rats. Mechanistically, our findings are consistent with αSYN oligomerization impairing vesicular DA handling and altering αSYN-associated vesicular machinery, changes that may increase free cytosolic DA and make it more available for oxidation into NM (Fig. 8).

**Figure 8.**
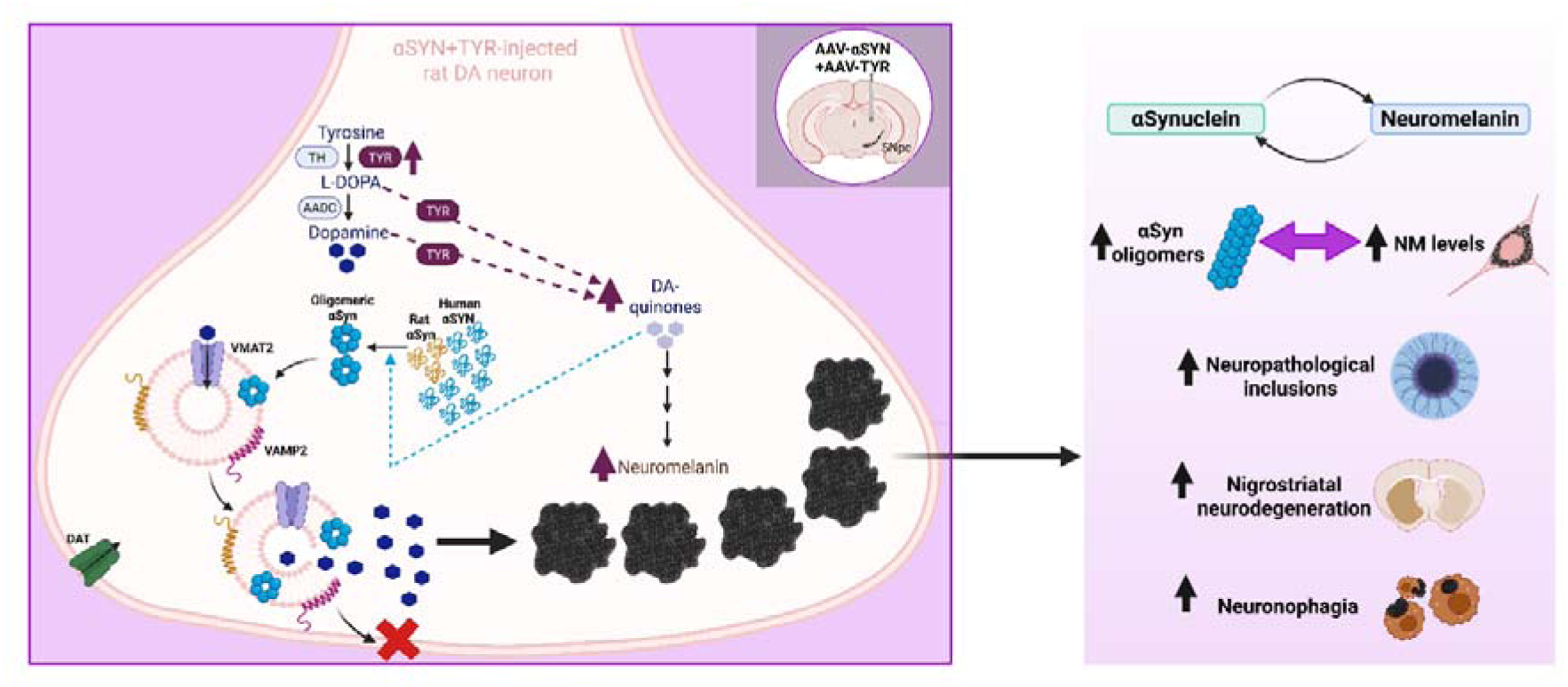
Schematic summary of the proposed αSYN-NM pathological feedback loop. Drawing was created with Biorender.com.

Our results extend earlier observations that αSYN and NM are spatially and biochemically linked in human PD brains. Previous studies showed that αSYN redistributes to NM granules early in PD and that NM granules can entrap aggregated αSYN (16,17). In particular, the lipid droplets present in NM granules are a nidus for αSYN accumulation (16), which may lead to LB formation. In line with this, experimentally induced NM in rodents and non-human primates stimulates the aggregation of endogenous αSYN into LB-like inclusions (12,23,25). While the exact mechanism of LB pathogenesis in pigmented neurons remains to be clarified, a maturation scheme has been proposed in human melanized nigral neurons, in which LB formation begins in preclinical PD, with filamentous αSYN-positive intracytoplasmic networks as the first signs of pathology (49). Consistent with this, we found that experimentally-induced NM accumulation in αSYN-overexpressing rats accelerated αSYN oligomerization and promoted sustained LB-like inclusion formation, which may render these neurons more susceptible to degeneration.

Besides direct NM-αSYN binding, in which αSYN is cross-linked to the pigment (17), other processes associated with NM formation and accumulation may also contribute to NM-induced αSYN aggregation. The synthesis of NM involves DA oxidation, a process that generates oxidative DA species, such as quinones, which act as NM precursors and can promote αSYN misfolding (50–52). Thus, NM accumulation and αSYN pathology may both arise, at least in part, downstream of a shared background of chronic oxidative/dopaminergic stress. In this context, high intracellular NM content may reflect the cumulative cytosolic redox burden experienced by these neurons, while progressively accumulated NM may itself contribute to cellular dysfunction. In parallel, the elevated SNCA expression observed in the NM-enriched human SNv may partly reflect this shared upstream environment rather than an entirely independent determinant of vulnerability. Reactive metals such as iron, which are bound to NM, may further amplify this process by catalyzing DA oxidation and promoting αSYN oligomerization (7,53–57).

In addition, intracellular crowding caused by progressive NM accumulation leads to a general proteostasis failure, accompanied by lysosomal and proteasomal deficits (11,12,58), which may further contribute to inclusion formation. Consistent with this, NM-laden cells exhibit a marked accumulation of p62-positive aggregates, which typically arise following inhibition of autophagy or proteasome-mediated intracellular degradation and are common components of LBs (15,59). Together, these pathological changes may contribute to the vulnerability of melanized nigral neurons in PD. Interestingly, we previously found that genetic ablation of αSYN in NM-producing, TYR-expressing mice does not reduce the formation of p62- and ubiquitin-positive LB-like inclusions (albeit lacking αSYN), nor does it prevent neurodegeneration (12). Together with the findings reported here, these results suggest that while αSYN may be dispensable for the initial formation of LBs and neurodegeneration, its pathological accumulation or elevated levels, as observed in PD patients, can accelerate NM-linked PD pathology.

Highly melanized neurons from both PD patients and NM-producing rats frequently show reduced intracellular TH levels (12). Brains from aged individuals also commonly exhibit TH downregulation within NM-laden SNpc neurons, in parallel with age-dependent αSYN accumulation (19,60). Although the functional significance of TH downregulation within melanized neurons remains unclear, diminished TH expression may predispose neurons to degeneration or may represent early stages of neuronal dysfunction/degeneration, even in the absence of overt PD. Indeed, while only a subset of individuals ultimately develops PD (1-5% of the population over 60, increasing with age (61), parkinsonian signs such as bradykinesia, stooped posture and gait disturbance are common among elderly individuals without clinical PD, reaching a prevalence of up to 50% in those older than 80 years (62,63). Notably, the TH-low, NM-rich (THNM□) neuronal population appears to be the most vulnerable to degeneration in PD and in TYR-expressing animals, as shown here. Although SNpc neurons with high NM content and reduced TH levels in aged human controls appear to maintain function based on PET and other neuroimaging measures, in NM-producing rodents TH downregulation is associated with impaired DA release and early motor and non-motor deficits that precede overt neurodegeneration (12,24,25,64). Similarly, Ndufs2-deficient mice with impaired mitochondrial respiration exhibit TH downregulation that coincides with the onset of motor deficits and levodopa-responsive parkinsonism, but precedes overt dopaminergic neuron loss (65). Boosting mesencephalic DA levels by overexpressing aromatic-l-amino acid decarboxylase (AADC) reverses these motor deficits (66). Likewise, optogenetic inhibition of DA neurons in rats induces TH downregulation and parkinsonism, whereas optogenetic reactivation restores TH expression and dopaminergic function (67,68). Together, these results point to TH downregulation as an early sign of neuronal dysfunction preceding neurodegeneration. The occurrence of neuronal dysfunction prior to cell death may have important therapeutic implications, as it provides a therapeutic window during which neuronal function could potentially be restored before irreversible degeneration. In the context of NM-linked neurodegeneration, reducing intracellular NM accumulation in AAV-TYR-injected rats, either by enhancing NM cytosolic clearance through the autophagy activator TFEB or by delaying age-dependent NM formation via VMAT2-mediated enhancement of dopamine vesicular sequestration, restores TH expression and markedly attenuates the neurodegenerative phenotype at both behavioral and neuropathological levels (12,24).

Our study has limitations inherent to the viral TYR model. Because NM synthesis is not a physiological feature of rodent neurons, ectopic TYR expression may not fully reproduce the lifelong kinetics and cellular adaptations associated with NM accumulation in the human SN. In addition, co-administration of two viral vectors could theoretically increase nonspecific effects of forced protein expression. However, dual-vector administration did not increase αSYN levels or the number of TYR-positive neurons compared with the corresponding single-vector groups, arguing against greater transgene burden as the main explanation for the αSYN+TYR phenotype. CMV promoter activity can also be enhanced by cellular stress (69), and although comparable numbers of TYR-positive SNpc neurons in TYR and αSYN+TYR animals at 2 months argue against a major effect, stress-dependent modulation of TYR expression per neuron over time cannot be excluded. Finally, the promoters used are not dopaminergic-cell-specific. Nevertheless, viral transduction and NM production are not equivalent: in this model, NM formation is selectively observed in nigral dopaminergic neurons, as pigment formation requires both TYR expression and the availability of cytosolic catecholamines for oxidation, whereas neighboring non-dopaminergic and glial cells expressing TYR do not accumulate NM or develop the associated pathology (12). Thus, the neuronal selectivity of NM formation and pathology argues against nonspecific effects of viral transduction or TYR expression per se.

Overall, this study identifies a previously underappreciated mechanism linking two hallmark PD features, such as αSYN and NM, and underscores the importance of incorporating NM into experimental models and clinical research to more accurately capture PD pathobiology. The reciprocal pathogenic relationship between αSYN and NM provides a mechanistic explanation for why highly pigmented neurons are especially vulnerable in PD. The ventral SN has the highest densities of NM-laden neurons (70,71) and these highly melanized cells are the first to degenerate in PD (30,60,71). Other, less pigmented cells are preserved initially but degenerate with disease progression (60). This suggests that the amount of intracellular NM relates to progressive PD pathogenesis, as shown in NM-producing animal and cellular models (12,23–25). Our findings recast the relationship between αSYN and NM as an active, self-reinforcing pathogenic loop rather than a passive coexistence. αSYN-driven vesicular dysfunction promotes NM synthesis, and excessive NM accumulation in turn enhances αSYN misfolding and inclusion formation, all of which create a cycle that accelerates neurodegeneration. Consequently, therapeutically disrupting this self-reinforcing αSYN-NM pathogenic cycle may offer a rational strategy to slow neurodegeneration and alter the course of disease progression.

## Acknowledgments

The authors acknowledge the technical and scientific assistance of Jordi Romero-Giménez, Sydney Microscopy & Microanalysis, the University of Sydney node of Microscopy Australia. We would also like to thank Heidi Cartwright for helping generate some of the figures.

## Declarations

### Ethical approval

All animal procedures were conducted in strict accordance with the European Directive (2010/63/UE) and Spanish regulations (Real Decreto 53/2013; Generalitat de Catalunya Decret 214/97) regarding the protection of animals used for experimental and other scientific purposes. Protocols were approved by the VHIR Ethical Experimentation Committee and the Generalitat de Catalunya (Protocol 11442). All human brains used in this study were collected by the Sydney Brain Bank following informed consent. The study was approved by the University of Sydney’s Human Research Ethics Committee (2021/HE000845).

### Data availability

Raw data on rodent and human dopamine neurons generated in this study is available upon request with Nanostring GeoMx Spatial Transcriptomic measurements available from https://doi.org/10.5281/zenodo.13626106 and https://doi.org/10.5281/zenodo.23390779. Human midbrain IHC for dopaminergic neurons in post-mortem tissue of control and PD patients https://doi.org/10.5281/zenodo.20278044. Neuromelanin in dopaminergic neurons in post-mortem tissue from control and Parkinson’s disease cases https://doi.org/10.5281/zenodo.20319696. All other data, code, protocols, and key laboratory materials used or generated in this study are documented in the Key Resource Table, together with their persistent identifiers, at https://doi.org/10.5281/zenodo.23337223.

### Author contributions

MV, ANV, YF and GMH conceived and designed the work; ANV, GR, JMCS, AP, TC, JM, AL, MGS and MV acquired and analysed the rodent data; YF, BH, AAM, FS, ZC and GMH acquired and analysed the human data; JH curated and managed the data; MV, ANV, MGS, YF, ZC and GMH interpreted the data and prepared the original draft of the manuscript; and all authors critically reviewed the manuscript and approved this version for publication.

